# A Cherry-Flavoured E-Cigarette Adduct, BPGA, Reprograms Alveolar Epithelial Cell Fate Through Epithelial-to-Mesenchymal Transition and Evasion of Apoptosis

**DOI:** 10.64898/2026.05.12.724520

**Authors:** Joseph Xavier, Yuefan Yu, Bhavani M Varma, Zhaolin Lu, Megha KB, Remya NS, Anil Kumar PR, Jorge Bernardino de la Serna

## Abstract

E-cigarettes have attracted significant attention as a safer substitute for conventional tobacco smoking. However, they have introduced new inhalable toxicants, including benzaldehyde-propylene glycol acetal (BPGA)—a chemical adduct produced by cherry-flavoured e-cigarettes. The health risks associated with such flavour-derived acetals remain insufficiently elucidated at the cellular level. This study investigated the role of BPGA in the progression of epithelial-to-mesenchymal transition (EMT)-like changes in alveolar epithelial cells (A549 cells).

A549 cells exposed to various concentrations of BPGA were analysed for cell viability, morphology, mitochondrial function, lysosomal health, and cytoskeletal integrity using viability assays and fluorescence imaging. Intracellular reactive oxygen species (ROS) production was quantified using the 2’,7’-dichlorodihydrofluorescein diacetate (DCFH-DA) assay. Antioxidant enzyme expression, inflammatory responses, and EMT-associated phenotypic alterations were evaluated using quantitative reverse transcription polymerase chain reaction (qRT-PCR) and immunofluorescence (IF) assays.

Exposure of alveolar epithelial cells to BPGA caused a concentration-dependent decrease in cell viability. BPGA exposure resulted in mitochondrial membrane depolarisation, lysosomal damage, cytoskeletal changes, and stress fibre formation, which altered cell morphology. It significantly increased intracellular ROS production. As a result, antioxidant enzyme levels were upregulated as a protective response. However, during severe oxidative stress, this response was overwhelmed. Excess ROS disrupted cellular homeostasis and initiated apoptosis, though not completely. ROS also acted as a signalling molecule, promoting the upregulation of inflammatory mediators. These changes were associated with altered EMT marker expression, suggesting that BPGA might drive EMT-like remodelling.

In conclusion, BPGA, a chemical adduct from e-cigarette vapour, induces alveolar injury by promoting oxidative stress, inflammation, and EMT-related changes, which may explain a mechanism by which e-cigarette exposure could lead to lung injury and pulmonary fibrosis.

Graphical abstract

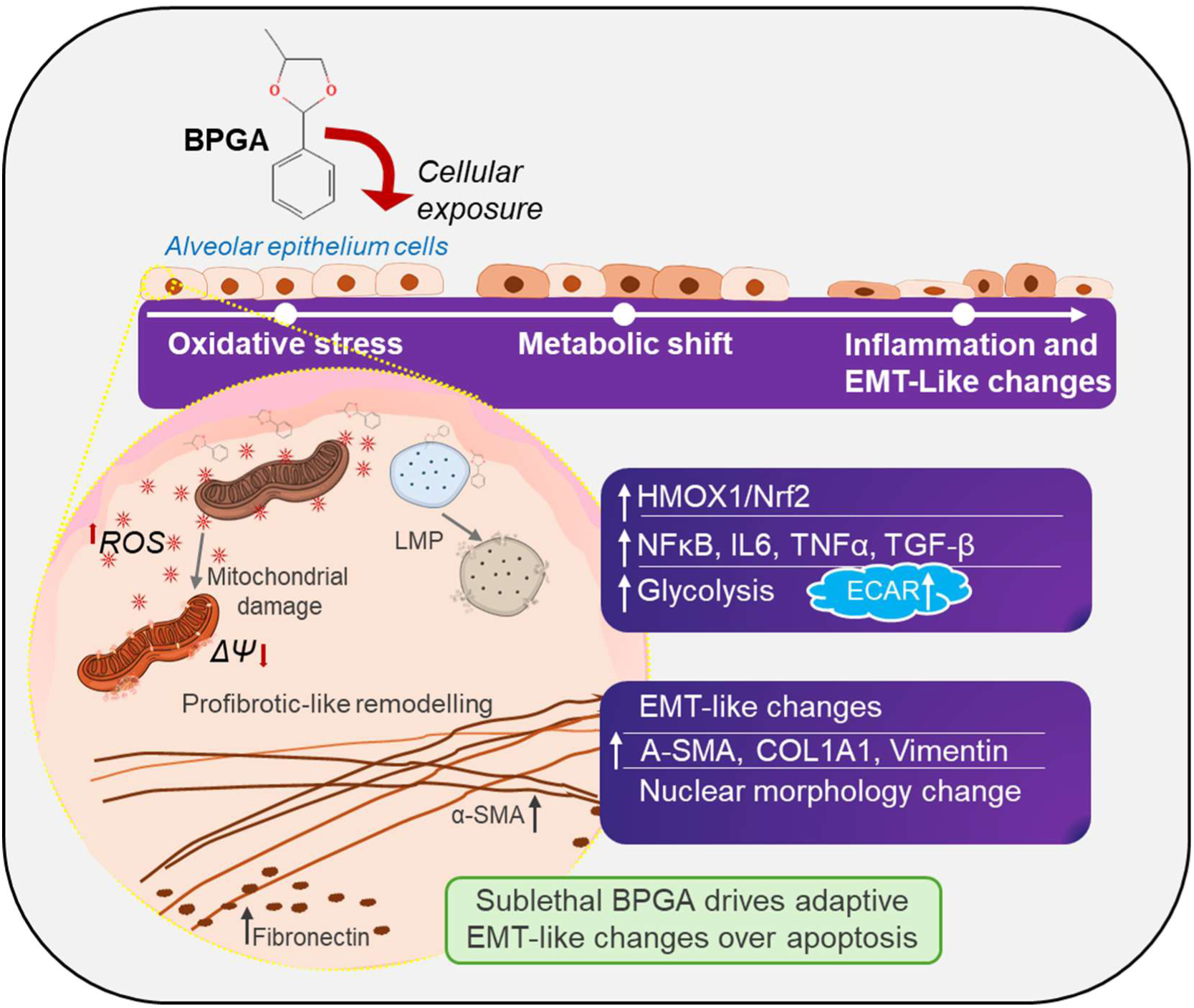

## 1. Introduction

E-cigarettes (e-cigs) are often seen and used as a way to help stop traditional smoking (Ali et al., 2023). It is important to emphasise that none of these devices is currently approved by any health institutions or organisations as therapeutic tools for treating health issues or addiction. Many users mistakenly believe these devices can help overcome addiction by offering a safer alternative (Sanchez-Rosario et al., 2024). Nevertheless, extensive research demonstrates that e-cigs also produce toxic substances upon thermal decomposition of e-liquids (Bekki et al., 2014; Margham et al., 2016). Despite these concerns, the use of e-cigs has increased across all age groups, with their global sales expanding rapidly. The total market value was approximately 37.96 billion USD in 2025 and is expected to reach around 200 billion USD by 2033 (Coherent Market Insights, 2025). The estimated number of active users worldwide approached 80 million in 2021. The prevalence of use among adults rose from 4.5% in earlier years to 6.5% in 2023 (Vahratian et al., 2025). Alarmingly, use among youth and adolescents has increased sharply in many countries, sometimes surpassing adult usage rates (Polosa et al., 2022).

E-cigs are battery-powered devices that generate an inhalable aerosol through the thermal vaporization of e-liquid, a formulation comprising flavoring agents, propylene glycol (PG), vegetable glycerin (VG), and optionally nicotine. The VG and PG act as humectants, helping to retain moisture and facilitate aerosol formation during heating. They also provide users with a distinctive “throat hit”. E-liquids often include cooling agents such as menthol, WS-3 (N-ethyl-p-menthane-3-carboxamide), and W-23 (2-isopropyl-N, 2,3-trimethylbutyramide) (Diomede, 2017), along with volatile organic compounds (VOCs), carbonyl compounds (such as acrolein, acetone, acetaldehyde, formaldehyde), metals, and inorganic compounds (including Cr, Ni, Mn, Pb, Al, Zn) (Kopa-Stojak & Pawliczak, 2025).

Manufacturers offer a wide range of unique flavours, with over 7,000 reported recently, to cater to diverse consumer preferences and boost user engagement (CDC, 2025). Users can personalise the flavour and nicotine strength of these products to suit their own tastes. The wide variety of flavouring agents and their unregulated use raise serious safety concerns (Czoli et al., 2019), particularly regarding long-term exposure risks. For instance, diacetyl, a flavouring agent used to produce a buttery taste in the food industry (Kassem et al., 2024), is generally considered safe for oral consumption but can cause an irreversible lung disease, bronchiolitis obliterans, also known as popcorn lung, when inhaled (Effah et al., 2022). Among the various flavourings available in the market, fruit-flavoured e-cigs are the most popular, especially among youth and adolescents. Benzaldehyde is the aromatic aldehyde used in e-cig liquids (E-liquids) to impart cherry flavours and is safe to ingest as a food additive, according to the “Flavour and Extract Manufacturers Association” (FEMA) and the US FDA. Nonetheless, safety guidelines may differ in cases of vaping-related exposure to benzaldehyde and its by-products. Notably, some e-cig cartridges contain up to 21 mg/mL of benzaldehyde as a flavourant (Tierney et al., 2016a). Although various studies have evaluated the toxicity of primary e-liquid components (Hua et al., 2019; Lerner et al., 2015), a comprehensive evaluation of the toxicity of secondary products from e-cigs is still lacking. Benzaldehyde can undergo an acetalization reaction with PG in the e-cig mixture under high temperatures during vaping, producing benzaldehyde propyl glycol acetal (BPGA), a secondary chemical adduct (Kerber & Peyton, 2022). Research indicates that 15-30% of flavour aldehydes convert to the corresponding propylene glycol acetal, and at least 40% of benzaldehyde propylene glycol acetal can carry over to e-vapour (Erythropel et al., 2019). These vapour products can further deposit on the airway lining along with other chemicals and remain there for days (half-life ≈ 44 hours for BPGA), indicating that they are not hydrolysing instantly and may cause adverse biological effects (Erythropel et al., 2019).

The physicochemical properties and exposure behaviour of flavour–propylene glycol acetal adducts during vaping remain incompletely characterised (S. Wu et al., 2023). Modelling studies suggest that these acetals exhibit altered gas–particle partitioning compared with their parent compounds. This shift favours association with the aerosol and condensed phases, thereby modifying inhalation exposure during vaping and indicating a potential for persistence in indoor environments after active use (Yeh et al., 2022). Several studies have revealed that the apolar nature of most e-cig aerosols interferes with the highly hydrophobic lipids and proteins of the pulmonary surfactant monolayer (Liekkinen et al., 2020, 2023) and disrupts its biophysical function (Goros et al., 2023; Graham et al., 2022; Martin et al., 2024). Pulmonary surfactant is the first line of defence against inhaled aerosols before they reach the gas-exchange barrier and facilitates breathing, preventing atelectasis (Andreassen et al., 2010; Dushianthan et al., 2020). Once this line is surpassed, e-cig aerosols will reach the alveolar epithelium and induce toxicity. Damage to alveolar epithelial cells and their underlying basement membrane from minor injuries or allergic irritants disrupts the normal air-blood barrier (Bi et al., 2025). These damaged cells trigger the inflammatory response by releasing cytokines that mediate inflammation (Zheng et al., 2025). Recurrent micro-injuries lead to the activation and generation of inflammatory macrophages (Long et al., 2023) and to profibrotic macrophage polarisation (Jiang et al., 2024), major players in the induction of lung fibrosis. Continuous exposure to respiratory pollutants can damage the normal barrier and eventually contribute to fibrosis through complex mechanisms (Makena et al., 2023), such as immune response dysfunction, oxidative stress, and chronic inflammation (Mariscal-Aguilar et al., 2024). This study furnishes evidence regarding the potential toxicity of benzaldehyde-propylene glycol acetal, a key flavourant adduct generated in a berry-flavoured e-cigarette formulations-towards alveolar epithelial cells.

## 2. Materials and methods

### 2.1. Cell culture maintenance

The human alveolar basal epithelial adenocarcinoma cell line (A549) is utilised for in vitro studies. These cells are generally employed in drug development, toxicological research, and EMT-related investigations. The A549 cell line (NCCS, India) was cultured in MEM (Himedia, India) supplemented with 10% fetal bovine serum (Gibco, South America origin, USA) and 1x antibiotic-antimycotic solution (Himedia, India), and subcultured every three days to reach 90% confluency. Cultures were incubated at 37°C in a humidified 5% CO₂ atmosphere to preserve cellular viability and physiological relevance.

### 2.2. 3-(4,5-dimethylthiazol-2-yl)-2,5-diphenyltetrazoliumbromide (MTT) assay

A549 cells were harvested using 1× trypsin-EDTA (Himedia, India), seeded into 96-well plate (Corning Costar, USA) at a density of 1×10^4 cells/well and incubated overnight to facilitate adhesion. The following day, the spent media was aspirated from each well and replaced with MEM (Himedia, India) containing various concentrations of BPGA (TCI, UK) (1, 3, 6, 9, 12, and 15 mM). MEM without BPGA served as the negative control; 0.1% DMSO (Himedia, India) as the vehicle control; and 1% phenol (Sigma-Aldrich, USA) as the positive control. The cells were subsequently incubated for 24 hours at 37°C in a humidified 5% CO₂ atmosphere. After incubation, the treatment media were removed and replaced with media containing MTT (0.5 mg/mL) (Himedia India), followed by incubation for an additional 3 hours at 37 °C with 5% CO₂. The spent medium containing MTT was aspirated, and DMSO was added to each well. The plate was placed in a shaking incubator in the dark for 30 minutes to solubilize the formazan crystals. Absorbance was measured at 570 nm using a microplate spectrophotometer (Multiskan SkyHigh, Thermo Fisher Scientific, USA). Cell viability, expressed as a percentage relative to the negative control, was plotted for each treatment group.

### 2.3. Neutral red uptake assay

The cell viability assay was conducted based on lysosomal uptake of neutral red. Briefly, A549 cells were harvested using 1× trypsin-EDTA and dispensed into 96-well plates (10⁴ cells/well), with overnight incubation to promote cell adhesion. Afterwards, the spent cell culture media was replaced with media containing different concentrations of BPGA (1, 3, 6, 9, 12, and 15 mM), 1% phenol as a positive control, 0.1% DMSO served as the vehicle control, with untreated cells designated as the negative control. The cells were incubated for 24 hours at 37°C with 5% CO₂. After incubation, the media was removed and replaced with neutral red staining solution (40 µg/mL) (SRL Chemicals, India), followed by an additional 3 hours of incubation under the same conditions. The staining solution was then aspirated, followed by a single wash with 1× PBS to remove unbound dye. Subsequently, neutral red dye was extracted using a destaining solution comprising 50% ethanol, 49% deionized water, and 1% glacial acetic acid. Absorbance was measured at 540 nm using a microplate spectrophotometer (Multiskan SkyHigh, Thermo Fisher Scientific, USA). The percentage of cell viability relative to the negative control was calculated and plotted for each treatment group.

### 2.4. Phase contrast microscopic imaging

A549 cells were seeded in a 24-well plate at a density of 5 × 10^4 cells/well and incubated overnight at 37°C in 5% CO₂ to facilitate attachment. Following incubation, the conditioned medium was aspirated and replenished with fresh medium supplemented with different BPGA concentrations (1, 3, 6, 9, 12, and 15 mM). Treatment without BPGA served as the negative control, and 0.1% DMSO was used as the vehicle control. The cells were incubated for 24 hours at 37°C with 5% CO₂. They were then observed under an inverted microscope (Olympus, Japan) equipped with an 18-megapixel CMOS colour camera (SC180, Olympus, Japan), Images were captured using an Olympus CKX53 inverted microscope with a 20x objective to assess cell morphology.

### 2.5. 2′,7′-Dichlorodihydrofluorescein diacetate (DCFH-DA) assay

A549 cells were seeded in a 96-well plate at a density of 1 × 10^4 cells per well and allowed to attach overnight under standard conditions. The next day, cells were treated with 10 μM 2′,7′-DCFH-DA (Invitrogen, USA) dissolved in 1x PBS for 30 minutes at 37°C. After incubation, the DCFH-DA solution was removed and replaced with MEM containing various concentrations of BPGA (3, 6, 9, 12, and 15 mM). MEM without BPGA served as the negative control, while 0.1% DMSO acted as the vehicle control. Hydrogen peroxide (Merck, USA) (100 μM) was used as the positive control. The cells were then incubated for a further 4 hours at 37°C with 5% CO₂. Fluorescent intensity (bottom reading) was measured using a Tecan Infinite F Nano+ fluorescent microplate reader (Thermo Fisher Scientific, USA) with excitation/emission filter settings of 488(20)/530(25) nm, and the data were plotted for each treatment group. To evaluate the effect of ROS scavenging, cells were co-treated with BPGA and ascorbic acid (1 mM), and ROS generation was assessed using the same assay conditions.

### 2.6. Tetramethylrhodamine (TMRM) assay

A549 cells were seeded in a 96-well plate at a density of 1×10⁴ cells per well. The cells were incubated overnight to facilitate attachment. The next day, the spent media was removed and replaced with MEM containing various concentrations of BPGA (1, 3, 6, 9, 12, and 15 mM); media without BPGA served as the negative control, while 100 μM hydrogen peroxide was used as the positive control. The cells were incubated for 24 hours at 37°C with 5% CO₂. After treatment, the media were discarded, and 100 nM TMRM diluted in MEM was added, and the mixture was incubated for 10 minutes at 37°C. The staining solution was removed, and the wells were washed once with 1× PBS. Fluorescent intensity (bottom reading) was measured using a Tecan Infinite F Nano+ fluorescent microplate reader (Thermo Fisher Scientific, USA) with excitation/emission filters set at 535 (10) / 595 (35) nm, and the results were plotted for each treatment group. Images were captured using an Olympus IX83 confocal laser scanning microscope; oil immersion objective of magnification 60x; with an excitation/emission wavelength of 560/590 nm and analysed using Fiji (ImageJ; NIH, USA).

### 2.7. Seahorse XF Cell Mito Stress Test

The effect of BPGA exposure on mitochondrial health in A549 cells was examined using the Seahorse XF Cell Mitostress Test (Agilent Technologies, Santa Clara, CA). Briefly, A549 cells were seeded in an Agilent Seahorse XFp Cell Culture Miniplate at a density of 2×10⁴ cells per well and cultured until near confluence. For calibration, the Agilent Seahorse XFp Sensor Cartridge was hydrated with deionised water 24 hours prior to the assay in a non-CO₂ incubator at 37°C. On the day of the assay, the cell culture medium was replaced with pre-warmed Seahorse XF assay medium, and the sensor cartridge was equilibrated with Seahorse XF Calibrant solution. Both plates were incubated at 37°C in a non-CO₂ environment for approximately one hour before the run. Working solutions of assay compounds were prepared immediately before use in XF assay medium. The injection order and final concentrations were as follows: (a) Vehicle control (assay medium with 0.1% DMSO) or BPGA (9, 12, or 15 mM), (b) Oligomycin (1 µM) to inhibit ATP synthase and measure ATP-linked respiration, (c) carbonyl cyanide-p-trifluoromethoxyphenylhydrazone (FCCP) (0.75 µM) to uncouple oxidative phosphorylation and determine maximal respiration and spare respiratory capacity (SRC), (d) a mixture of Rotenone and antimycin A (1 µM) was used to inhibit Complex I and III, respectively, while measuring non-mitochondrial respiration. Oxygen consumption rate (OCR) data were normalised to basal OCR (mean of the first three measurements before the first injection). Changes in OCR (ΔOCR) were calculated per manufacturer recommendations to derive mitochondrial parameters. Extracellular acidification rate (ECAR) was also monitored and calculated as an indicator of glycolytic metabolism. Graphs and statistical analyses (ordinary two-way ANOVA) were generated using GraphPad Prism 9 (GraphPad Software, La Jolla, CA).

### 2.8. Lysosomal staining

Acridine orange (AO) staining was performed to visualise and analyse lysosomal integrity. Acridine orange, a weak base, enters the cell and accumulates in the acidic environment of the lysosome. In the cytosol, AO exists as monomers and emits green fluorescence. However, once it forms a dimer due to the acidic environment of the lysosome, it shows a metachromatic shift to red. Briefly, A549 cells were harvested and seeded into a 24-well plate containing sterile glass coverslips at a density of 5 × 10⁴ cells per well, then incubated overnight at 37°C with 5% CO₂ to allow for cell attachment. The spent media was removed and supplemented with fresh media containing increasing concentrations of BPGA (3, 6, and 9 mM); media alone served as the negative control, and 1% phenol was the positive control. The cells were incubated for 24 hours. The treatment media was replaced with 1x PBS containing 2.5 µg/mL of AO (Himedia, India) and incubated for 20 minutes. The cells were washed once with 1x PBS and images were captured using an Olympus IX83 confocal laser scanning microscope; oil immersion objective of magnification 60x with excitation/emission laser settings of 488/525 nm for AO monomers and 488/650 nm for AO dimers. Image analysis was performed using Fiji-ImageJ (NIH, USA). The mean fluorescent intensity (MFI) ratio of AO dimer to AO monomer was calculated and plotted for each treatment group.

### 2.9. 3D Holotomographic Microscopy

A total of 7.5 × 10⁵ A549 cells were seeded into 35 mm ibiTreat dishes (Ibidi, Germany) and cultured for 24 hours in complete cell culture medium supplemented with 10% serum and 1% penicillin-streptomycin-glutamine until reaching approximately 80% confluency. After incubation, the medium was replaced with fresh medium containing 9 mM or 12 mM BPGA, 0.1% DMSO as a vehicle control, media alone as a negative control, and TGF-β1 (20 ng/mL) as a positive control. The dishes were then transferred to the top stage incubator (Tokaihit, Japan), and holotomographic imaging was performed using a CX-A 3D Live Cell Explorer (Nanolive, Switzerland) under standard culture conditions (37°C, 5% CO₂). The sample holder was pre-equilibrated under standard conditions for at least 3 hours to minimise focus drift due to thermal instability. The imaging cycle was performed in 6-minute intervals for 200 cycles, resulting in a total acquisition period of 20 hours. The conditions were optimised to prevent medium evaporation and dehydration-induced cell death, and to enable comparison with previously obtained MTT data. Additionally, to capture cellular events within the 24–48-hour window, cells were maintained under the respective treatment conditions for 24 hours, then transferred to the sample holder, and image acquisition was performed using the aforementioned settings. Cell confluency, surface area, dry mass, and cellular granularity were quantified using Eve Analytics software (EA-UM-001 version 1.0, Nanolive, Switzerland).

In Nanolive experiments (using the 3D Cell Explorer platform, which measures the refractive index, RI, of live cells), these three parameters are defined as follows:

(a) Eccentricity: A shape descriptor of the segmented object (cell or organelle). It is derived from the best-fit ellipse to the 2D projection (or 3D volume) of the object and describes how elongated or irregular the shape is relative to a perfect circle/sphere. Values range from 0 (perfect circle) to 1 (a line). A higher eccentricity indicates a more elongated or asymmetric object. In Nanolive’s STEVE software, it is typically computed from the 2D projected mask of each segmented region.

(b) Dry Mass: The total amount of non-aqueous, organic material (proteins, lipids, nucleic acids, carbohydrates) within a segmented region, calculated directly from the refractive index using the Gladstone–Dale relationship:

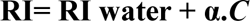

where ***C*** is the concentration of dry material (g/mL) and **α** is the specific refractive index increment (∼0.190 mL/g for most biological materials). Dry mass (in picograms, pg) is then:

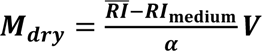

where V is the volume of the segmented region. It is a label-free, quantitative readout of cellular or organellar biomass.

(c) Granularity: A texture/heterogeneity descriptor that quantifies the spatial variation in refractive index within a segmented object. It reflects how non-uniform the internal RI distribution is — essentially, how “textured” or “grainy” the interior of the object appears in the RI tomogram. Mathematically it is typically expressed as the standard deviation (or variance) of the RI values across all voxels within the segmented region:

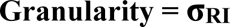

Higher granularity indicates greater internal heterogeneity (e.g., dense organelles, condensates, lipid droplets, or aggregates embedded in a lower-RI cytoplasm). It is particularly sensitive to sub-cellular compartmentalisation and phase-separated structures.

These three parameters together give a powerful, label-free characterisation of cellular state, particularly relevant for detecting condensate formation, organelle remodelling, or changes in biomass distribution.

### 2.10. Actin-filament staining

Actin filament (F-actin) staining was performed using rhodamine-conjugated phalloidin, which binds tightly to F-actin, preventing its depolymerisation and enabling the visualisation of the F-actin structure. A549 cells were harvested and seeded into µ-Slide 15-Well plates (Ibidi GmbH, Germany) at a density of 4,000 cells per well, then incubated overnight at 37°C with 5% CO₂ to allow for cell attachment. The spent culture media were subsequently replaced with media containing increasing concentrations of BPGA (3, 6, and 9 mM); media alone served as the negative control, and 20 ng/mL TGF-β served as the positive control. The cells were incubated for 24 hours at 37°C with 5% CO₂. After treatment, the media was removed, and the cells were washed once with 1x PBS. Cells were fixed in 4% paraformaldehyde (Sigma-Aldrich, USA) for 20 minutes at room temperature, then washed three times in 1x PBS for 5 minutes each. The cells were then permeabilised with 0.1% Triton X-100 for 5 minutes and washed thrice with 1x PBS for 5 minutes each. Rhodamine-phalloidin (Abcam, USA) (1× solution in 1% BSA in 1x PBS was added, and cells were incubated for 30 minutes at room temperature in the dark. After staining, the solution was removed, and cells were washed once with 1x PBS, then counterstained with 4′,6-diamidino-2-phenylindole (DAPI) (Molecular Probes, Life Technologies, USA) for 5 minutes. Finally, the cells were washed with 1x PBS, and antifading mounting medium was added to each well to preserve the staining and images were captured using an Olympus IX83 confocal laser scanning microscope; oil immersion objective of magnification 60x with excitation/emission settings of 405/461 nm for DAPI-stained nuclei and 560/590 nm for rhodamine–phalloidin–stained F-actin. Image analysis was performed using Fiji-ImageJ (NIH, USA).

### 2.11. Differential Interference Contrast (DIC) imaging

The morphology change upon BPGA treatment was visualised using DIC microscopy. Briefly, A549 cells were harvested, seeded onto sterile glass coverslips in a 24-well plate at a density of 5 × 10⁴ cells per well, and incubated overnight at 37°C with 5% CO₂ to allow cell attachment. The spent media was removed and supplemented with fresh media containing increasing concentrations of BPGA (3, 6, and 9 mM); media alone served as the negative control, and 20ng/mL TGF-β was used as positive control. The cells were incubated for 24 hours. After 24 hours of incubation, cells were imaged live using inverted microscope equipped with DIC optics with Nomarski prisms (Olympus, Japan) with a 60x oil-immersion objective using polarised light. The images were analysed using Fiji-imageJ (NIH, USA)

### 2.12. Cell cycle analysis using flow cytometry

To study the effect of BPGA treatment on the progression of the cell cycle, flow cytometry was used. A549 cells were harvested and seeded in a 6-well plate at a seeding density of 3 × 10⁵ cells per well and incubated overnight at 37°C with 5% CO₂ to allow cell attachment. The spent media were removed and replaced with fresh media containing increasing concentrations of BPGA (3, 6, and 9 mM); cells treated with media alone served as the negative control, and 20 µg/mL mitomycin was used as the positive control. The cells were incubated for 24 hours. After incubation, spent media were collected to recover any detached cells due to BPGA exposure, and adherent cells were harvested by trypsinisation. The collected cells from both the spent media and the attached cell group were pooled and subsequently fixed in 70% ethanol and stored overnight at −20°C. The cells were then pelleted at 4,000 RPM for 2 minutes. Cells were permeabilised with 0.25% Triton X-100 in PBS for 15 minutes on ice. The permeabilised cells were collected by centrifugation at 4,000 RPM for 2 minutes and incubated in PBS containing 10 µg/mL RNase A and 10 µg/mL PI for 30 minutes at room temperature in the dark. After incubation, the cells were pelleted, resuspended in PBS, strained through a 40 µm cell strainer, and loaded for flow cytometry. Singlet cells were gated using forward scatter – area (FSC-A) vs forward scatter – height (FSC-H) to exclude doublets, followed by side scatter – area (SSC-A) vs FSC-A for size and granularity, and a minimum of 10,000 cells from the singlet population were used for further analysis. The DNA histogram was then plotted and the cell cycle distribution (G0/G1, S, and G2/M phases) was determined using the Dean-Jett-Fox (DJF) model with a standard 1-cycle fit in FCS Express. (De Novo Software).

### 2.13. Comet assay

During apoptosis, DNA fragmentation often occurs. Therefore, BPGA-induced DNA fragmentation was analysed using the comet assay, also known as single-cell gel electrophoresis. In the comet assay, fragmented DNA migrates during electrophoresis and appears as comet tails.

Microscopic glass slides were coated with 1% normal agarose by dipping for 30 seconds; the back side of the slide was then wiped to remove excess agarose and dried overnight at room temperature with the agarose side facing upwards. A549 cells were harvested using trypsin-EDTA, seeded in a 6-well plate at a density of 3 × 10⁵ cells per well, and incubated overnight at 37°C with 5% CO₂ to allow attachment. The medium was then replaced with fresh medium containing increasing concentrations of BPGA (3, 6, 9, and 12 mM). Cells treated with medium alone served as the negative control. After 24 hours of incubation, detached cells were collected from the spent medium, while adherent cells were harvested by trypsinization. The collected cells from both the spent medium and the adherent group were pooled, centrifuged at 1000 RPM for 3 minutes, and resuspended in 1x PBS.

Low-melting-point agarose (1%) was prepared in a 60°C water bath. The cell suspension (40 µL) was mixed with low-melting-point agarose (140 µL), and 75 µL of the mixture was dropped onto the agarose-precoated glass slide and immediately covered with a coverslip. allowed to solidify for 5 minutes at 4°C in a refrigerator. After solidification, the coverslip were removed, and slides were incubated in pre-chilled lysis buffer (2.5 M sodium chloride, 100 mM disodium EDTA, 100 mM Tris-HCl, 200 mM sodium hydroxide, 10% dimethyl sulfoxide, and 1% Triton X-100; DMSO and Triton X-100 were freshly added; pH 10) for 1 hour.

Following lysis, the slides were transferred to an electrophoresis tank filled with alkaline buffer (1 mM Na₂EDTA, 300 mM NaOH, pH 13) and incubated for 40 minutes at 4°C to allow DNA unwinding. Electrophoresis was performed at 25 V, 300 mA for 20–30 minutes. After electrophoresis, the slides were washed with neutralising buffer (400 mM Tris-HCl, pH 7.5) for 10 minutes and rinsed with water. The cells were stained with 20 µg/mL ethidium bromide and imaged using an Olympus IX83 confocal laser scanning microscope using a 520/605 nm laser, with 20x objective. The images were imported into Fiji-ImageJ, and the Open Comet plugin was used to analyse the percentage of tail DNA. The tail DNA percentage was then plotted against BPGA treatment.

### 2.14. Acridine orange (AO)/ethidium bromide (EB) assay

Significant increases in ROS may trigger apoptosis in epithelial cells. To check whether BPGA treatment induces apoptosis in A549 cells, an AO/EB assay was performed. A549 cells were harvested using trypsin-EDTA, seeded in a 6-well plate at a density of 3 × 10⁵ cells per well, and incubated overnight at 37 °C with 5% CO₂ to allow attachment. The medium was then replaced with fresh medium containing increasing concentrations of BPGA (3, 6, 9, and 12 mM). Cells treated with medium alone served as the negative control, and cells treated with 40 µM cisplatin served as the positive control. After 24 hours of incubation, detached cells were collected from the spent medium, while adherent cells were harvested by trypsinisation. The collected cells from both the spent medium and the adherent group were pooled, centrifuged at 1000 RPM for 3 minutes, and resuspended in ice-cold 1x PBS (25 µL). Subsequently, 2 µL of AO/EB solution (AO: 100 µg/mL, EB: 100 µg/mL in PBS) was added. A 10 µL aliquot of the stained cell suspension was loaded onto a clean glass slide and secured with a coverslip. The cells were then visualised using an Olympus IX83 confocal laser scanning microscope; objective 60x oil immersion, using 488 nm and 576 nm lasers. The images were analysed using Fiji ImageJ (NIH, USA). A custom macro code was generated to separate and analyse the green-only ROI (normal cells), red-only ROI (necrotic cells), and red–green overlap ROI (apoptotic cells). The apoptotic index was calculated by dividing the number of apoptotic cells by the total number of cells and converting the result into a percentage.

### 2.15. RNA isolation and cDNA synthesis

The extraction of RNA was performed using the HiPura total RNA miniprep purification kit (Himedia, India) according to the manufacturer’s standard protocols. Briefly, A549 cells were harvested and seeded into 60 mm cell culture dishes at a seeding density of 5 × 10⁵ cells per well and allowed to adhere overnight at 37°C in a humidified atmosphere with 5% CO₂. The following day, the spent media were aspirated and replaced with fresh media containing BPGA at increasing concentrations (3, 6, and 9 mM), with untreated media as the negative control. Cells were incubated for 24 hours under the same conditions. Post-treatment spent media were collected to recover any detached cells due to BPGA exposure, and adherent cells were harvested by trypsinisation. Pelleted cells from both the spent media and the corresponding adherent cell fraction were lysed in 600 µL of the kit’s lysis buffer by vortexing for 15 seconds at room temperature. Next, the lysates were passed through HiShredder at 12,000 RPM for 2 minutes to remove insoluble debris. An equal volume of 70% ethanol was added to the lysate and mixed thoroughly by pipetting. The resulting mixture was loaded onto the miniprep spin column and centrifuged at 10,000 RPM for 15 seconds to allow RNA to bind to the membrane. The membrane-bound RNA was sequentially washed with 700 µL of RWI buffer and 500 µL of WS buffer by centrifugation at 10,000 RPM for 1 minute each. The membrane was dried by a final wash with 500 µL of WS buffer and centrifuged at 10,000 RPM for 2 minutes. Subsequently, the total RNA was eluted with 50 µL of elution buffer into a fresh-capped collection tube by centrifugation at 10,000 RPM for 1 minute. The yield and purity of the eluted total RNA were assessed using a Nanodrop spectrophotometer (Thermo Fisher Scientific, USA), and the samples were stored at −80°C until further analysis.

cDNA synthesis was performed using PrimeScript™ RT Master Mix (RR036A, Takara, Japan) according to the manufacturer’s protocol. Briefly, a 20 µL reaction mixture was prepared by mixing 4 µL of 5X PrimeScript RT Master Mix, 1 µg RNA template, and RNase-free water to a final volume of 20 µL. A two-step reverse transcription (RT) protocol consisting of an RT reaction at 37°C for 15 minutes followed by heat inactivation at 85°C for 5 seconds was performed on the CFX96 real-time PCR system (Bio-Rad, USA). The resulting cDNA was diluted using the Easy Dilution Buffer supplied with the kit for downstream applications.

### 2.16. Quantitative Polymerase Chain Reaction (qPCR)

The fold change of various genes related to antioxidant response, apoptotic pathway, inflammatory response, and epithelial to mesenchymal transition was calculated using iTaq Universal SYBR Green Premix (1725121, Bio-Rad, USA) following the manufacturer’s standard protocol. A 10 µL reaction mixture was prepared by combining 5 µL of 2X master mix, 500 nM of each forward and reverse primer, 10 ng of cDNA template, and nuclease-free water (MB064, Himedia) to reach a final volume of 10 µL. The mixture was thoroughly mixed, loaded onto a skirted 96-well PCR plate (Bio-Rad, USA), and sealed with an optically transparent film. The plates were briefly centrifuged to eliminate any air bubbles. Amplification was performed using a two-step thermal cycling program on a CFX96 thermocycler (Bio-Rad, USA), followed by melt curve analysis to confirm product specificity. Ct values were recorded, and relative fold change was calculated as 2^(-ΔΔCt), normalised to either GAPDH or β-Actin as an internal control. The nucleotide sequences of the primer pairs used for the qPCR analysis of various genes are listed in supplementary table 1.

### 2.17. Immunofluorescence assay

Immunofluorescence stains were performed to examine protein expression and localisation. We carefully adapted previous protocols (García et al., 2016; Stanly et al., 2016) to avoid artefactual cluster generation for precise quantification. A549 cells were harvested and seeded into µ-Slide 15-Well plates (Ibidi GmbH, Germany) at a density of 4,000 cells per well, followed by overnight incubation under standard conditions to allow cell attachment. The culture medium was subsequently replaced with complete culture medium containing increasing concentrations of BPGA (3, 6, 9 mM). Media alone served as the negative control, while 25 µM Actinomycin D was used as the positive control for HO-1/HMOX-1 and cleaved caspase-3, and 20 ng/µL TGF-β1 was used as the positive control for EMT-related protein markers. Cells were incubated under these conditions for 24 hours. After treatment, the media were removed, and cells were washed once with 1× PBS, then fixed with 4% paraformaldehyde for 20 minutes at room temperature. After fixation, cells were washed three times with 1× PBS (5 minutes each). Permeabilisation was performed with 0.25% Triton X-100 in 1× PBS for 5 minutes, followed by three washes with 1× PBS (5 minutes each). Blocking was performed in 1% BSA in PBST containing 300 mM glycine for 1 hour at room temperature. Subsequently, cells were incubated overnight at 4°C with primary antibodies (listed in Table 1) diluted in antibody dilution buffer (1% BSA in PBST). The following day, the primary antibody solution was removed, and cells were washed three times with 1× PBS (5 minutes each). Fluorophore-labelled secondary antibodies (diluted in antibody dilution buffer) were added and incubated for one hour at room temperature. Cells were washed thrice with 1× PBS and counterstained with DAPI (Molecular Probes, Life Technologies, USA) for 5 minutes at room temperature. Finally, cells were washed three times with 1× PBS, mounted in antifade mounting medium (G Biosciences, USA), and imaged using a confocal laser scanning microscope (Olympus IX83, Japan) with a 60x oil-immersion objective. Imaging was performed with the following laser settings: 485/530 nm for excitation/emission of the Alexa Fluor 488-labelled secondary antibody and 405/461 nm for DAPI-stained nuclei. Image analysis was performed using ImageJ FIJI (NIH, USA). The integrated density of each image was quantified and normalised to the number of cells determined by nucleus counts. Fold change was calculated by dividing the normalised integrated density of each image by the average normalised integrated density of the negative control. The nuclear aspect ratio and roundness were obtained from DAPI-stained nuclei using the particle analysis workflow in Fiji ImageJ.

**Table 1.** Details of antibodies used for the immunofluorescence assay.

| Antibody | Catalogue number and make | Dilution |
| --- | --- | --- |
| Heme Oxygenase 1 (HO-1/HMOX1) Rabbit pAb | A21452, Abclonal | 1:50 |
| α-Smooth Muscle Actin (ACTA2) Rabbit pAb | A7248, Abclonal | 1:100 |
| Fibronectin (FN) Rabbit pAb | A12932, Abclonal | 1:100 |
| Cleaved Caspase-3 (Asp175) (5A1E) Rabbit mAb | 9664, Cell Signalling technology | 1:400 |
| Goat pAb to Rb IgG Alexa Fluor 488 | ab150077, Abcam | 2 µg/mL |

For α-SMA Ripley’s L(r)-r: spatial organisation comparison and nearest neighbour analysis (Kiskowski et al., 2009) confocal fluorescence images were acquired using a 60x oil immersion objective, laser settings: detector voltages-600, Laser LD405 transmissivity-0.90, Laser LD488 transmissivity-0.5, Objective Lens na-1.42. Image stacks were analysed using ThunderSTORM plugin in Fiji. Images were pre-processed with a B-spline wavelet filter (scale=2, order=3) to enhance the signal-to-noise ratio and suppress background noise. The fluorophore centroids were detected as local maxima in the filtered images, using 8-neighbourhood connectivity and a threshold defined as the standard deviation of the first wavelet component. The localisation was achieved by integrated Gaussian PSF with weighted least squares, including multi-emitter fitting to account for crowded field. Subsequently, the co-ordinates of the centroids were analysed in R (version 4.x) using the *spatstat* and *ggplot2* packages. Each cell was represented as a point pattern object (ppo) with margins defined by the bounding box of the data. For each cell, the nearest-neighbour distance (NND) was calculated using the *nndist()* function and mean ± SD values were recorded. Spatial clusterinwas further evaluated using Ripley’s L(r)–r function, computed with *Lest(L-function estimate)* over distances from 0–20 µm at 0.1 µm intervals. Positive deviations of L(r)–r above zero indicated spatial clustering, while negative values suggested dispersion relative to complete spatial randomness. Individual Ripley plots were also produced for each cell to illustrate heterogeneity in bundle organisation.

### 2.18. Statistical analysis

All experiments were conducted in triplicate. A standard one-way ANOVA, followed by Dunnett’s multiple comparison test (unless otherwise specified), was used to identify significant differences relative to the control. For image analysis, fluorescent images from at least five different locations were captured for each treatment group, and the integrated density was quantified and normalised by the number of nuclei. Data are shown as mean ± standard deviation (SD). Statistical significance was indicated as *p < 0.05, *p < 0.01, ***p < 0.001.

## 3. Results

### 3.1. BPGA induces a concentration-dependent cytotoxicity

BPGA exhibited a concentration-dependent cytotoxicity in A549 cells, as demonstrated by both quantitative (Figure 1A & B) and morphological (Figure 1C) cell viability assessments. A significant reduction in cell viability was observed from 6 mM onwards, with a rapid decline evident even after short exposure periods, such as 2 and 4 hours (Figure S1). Increasing concentrations of BPGA caused progressive disruptions in cellular morphology. Treatment with BPGA resulted in distorted cell morphology, a notable increase in detached cells, and elongated cells compared with control groups (Figure 1C).

**Figure 1.**
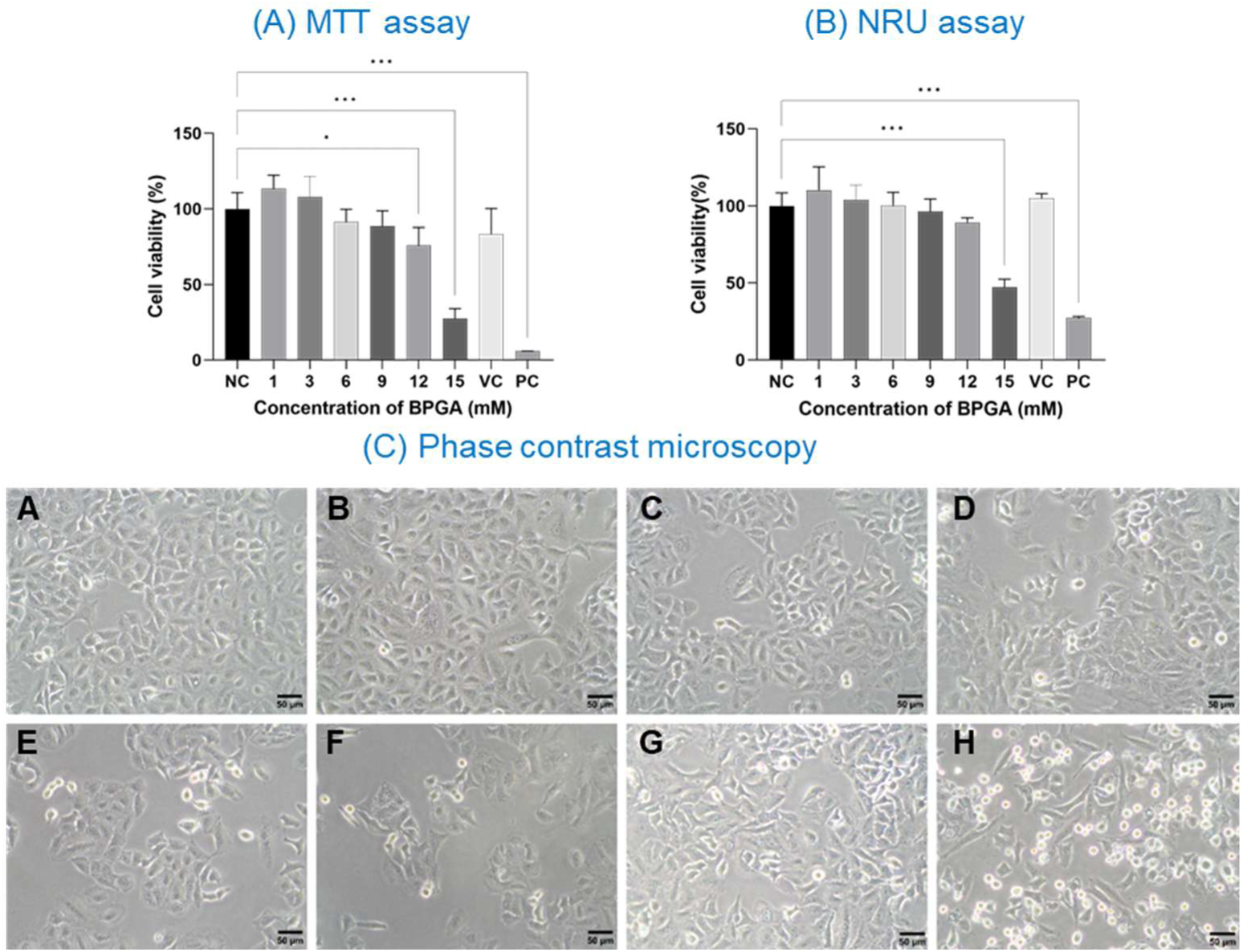
(A) Cell viability is expressed by the ability of A549 cells to reduce MTT into formazan crystals. A549 Cells (1 × 104 cells/well) were exposed to BPGA (1mM, 3mM, 6mM, 12mM, 15mM), 0.1% DMSO (VC) and 1% Phenol (PC) for 24 h. (B)Cell viability is expressed by the lysosomal activity of the A549 cells. A549 Cells (1 × 104 cells/well) were exposed to BPGA (1mM, 3mM, 6mM, 12mM, 15mM), 0.1% DMSO (VC) and 1% Phenol (PC) for 24 h. Data are presented as mean ± standard deviation. Statistical comparisons were performed using the ordinary one-way ANOVA test, followed by Dunnett’s multiple comparison test; *p < 0.05, **p < 0.01, ***p < 0.001. Mitochondrial depolarisation induced by BPGA treatment. (C) Phase contrast images showing the morphology of A549 cells treated with various concentrations of BPGA in 0.1% DMSO; (A) NC, (B) VC, (C) 1 mM, (D) 3 mM, (E) 6 mM, (F) 9 mM, (G) 12 mM, and (H) 15 mM; the scale bar represents 50µm.

Quantitative cell viability analysis using the MTT assay (Figure 1A) demonstrated that exposure to 1 and 3 mM BPGA resulted in mean cell viabilities of 105±6.4 % and 97.4±6.8%, respectively, indicating no significant cell death compared to controls. Cell viability decreased at 6 mM (84.6±6.8%), 9 mM (81.6±8.7%), 12 mM (67.3±5.0%), and 15 mM (23.5±3%).

Statistically significant reductions in viability were observed at exposures ≥12 mM (ANOVA, p < 0.001). The IC50 was determined as 13.5 mM (95% CI 12.8–14.3 mM; Hill slope −8.4), calculated by nonlinear regression using a log(inhibitor) versus normalised response, variable-slope model. Acute BPGA exposure (2–4 hours) also caused rapid and substantial declines in cell viability at higher concentrations (≥15mM) (S1 A&B), emphasising BPGA’s swift and potent cytotoxic effects. The neutral red uptake (NRU) assay demonstrated comparable trends in cell viability alterations following BPGA treatment, consistent with MTT assay results (Figure 1B). The NRU assay, which assesses cell viability based on lysosomal health, indicated that BPGA treatment at up to 9 mM had minimal effects, with mean viability values of 101±11.4 (1 mM), 97.8±9.5 (3 mM), 95.1±9.0 (6 mM), and 90.3±6.4 (9 mM). However, at 12 mM and 15 mM, mean cell viability decreased to 85.4±3.0 and 44.3±5.0, respectively.

### 3.2. BPGA exposure increases intracellular ROS generation and modulates the antioxidant response in A549 cells

BPGA exposure on A549 cells resulted in a significant concentration-dependent elevation in intracellular reactive oxygen species production, as determined by the DCFH-DA assay (Figure 2B). The relative fluorescent intensity (RFU) increased progressively with increased BPGA concentrations. Lower concentrations of BPGA (≤3 mM) did not result in a statistically significant generation of intracellular ROS in comparison with negative control (untreated); however, BPGA concentrations above 6 mM led to a significantly elevated fluorescence, indicating increased ROS generation (ANOVA, p < 0.01 to p < 0.001). The mean fluorescent intensity (MFI) (a.u.) increased from 4765±547 to 8237±1415 (3 mM), 10857±1624 (6 mM), 12309±704 (9 mM), 13303±1693 (12 mM) and 18051±5093 (15 mM). These results demonstrate that BPGA at concentrations of 6 mM or higher induces robust oxidative stress in A549 cells, suggesting its potential to disrupt redox homeostasis by increasing intracellular ROS levels.

**Figure 2.**
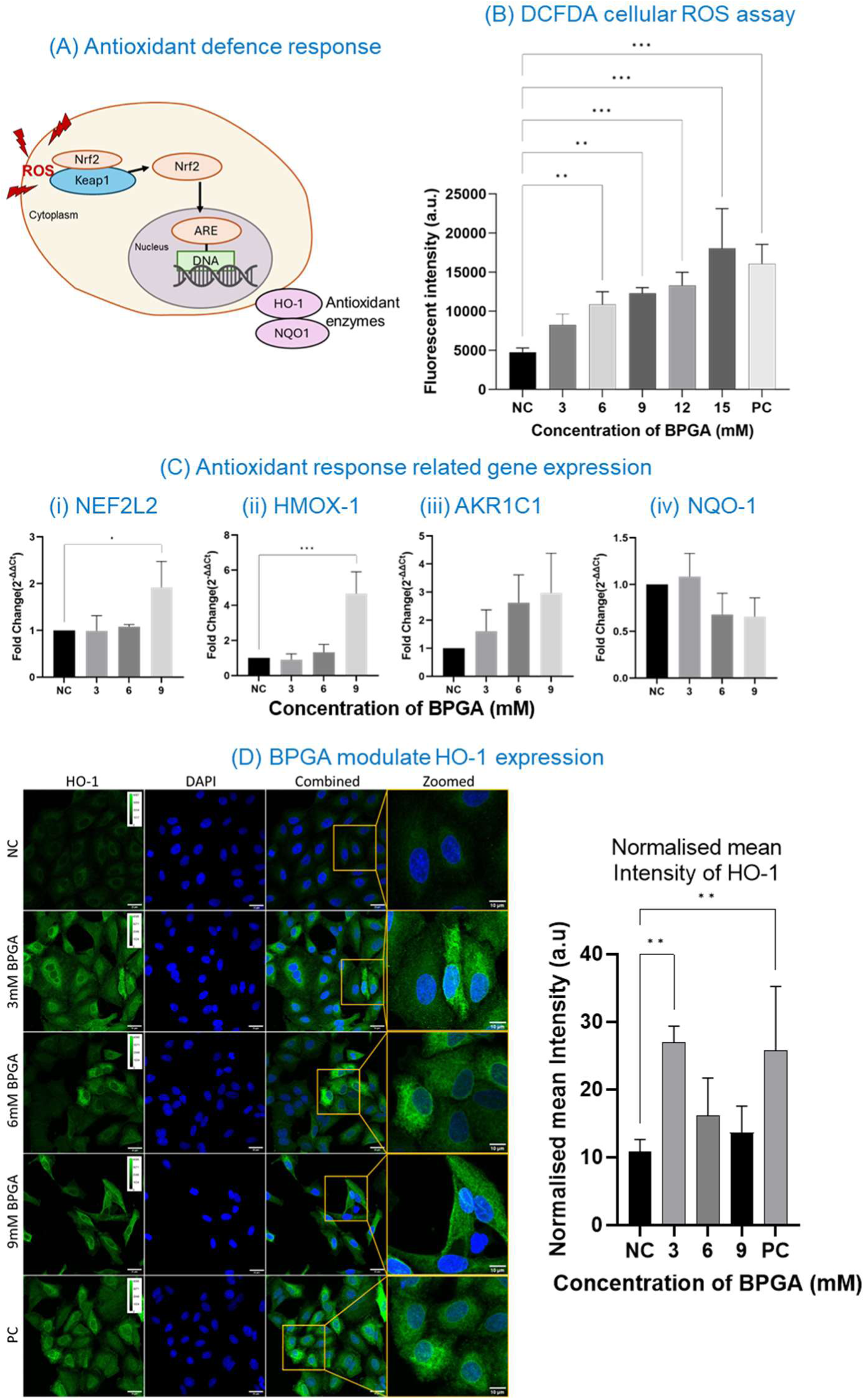
BPGA-induced antioxidant response (A) Antioxidant response pathway schematic, (B) Cells were treated with various concentrations of BPGA (3, 6, 9, 12, 15 mM), 100µM H₂O₂ (PC) for 4h and ROS generation was assessed using DCF-DH assay, (C) Quantitative PCR analysis of antioxidant response-related genes after BPGA treatment, normalised with housekeeping genes. (i) NFE2L2, (ii) HMOX-1, (iii) AKRIC1, (iv) NQO-1, (D) HO-1 expression in A549 cells treated with various concentrations of BPGA, 25µM Actinomycin A (PC) for 24h. scale bar represents 25µm (10µm for zoomed images).

The significant increase in ROS can further activate the antioxidant defence response as a compensatory mechanism (Figure 2A). During the normal ROS-mediated stress response, elevated ROS can activate the Nrf2–Keap1 pathway by modifying a cysteine residue in Keap1, thereby reducing its ability to ubiquitinate Nrf2. As a result, Nrf2 accumulates, translocate to the nucleus and binds to the antioxidant response element (ARE) in DNA, thereby upregulating genes encoding various antioxidant enzymes (Qu et al., 2023).

Therefore, the changes in antioxidant gene expression upon exposure to BPGA were evaluated using quantitative PCR, and fold changes (2^-ΔΔCq^) of nuclear factor erythroid 2-related factor 2 (NFE2L2) (gene encoding the transcription factor Nrf2), Heme oxygenase-1 (HMOX-1) (gene encoding heme oxygenase 1), Aldo-Keto Reductase Family 1 Member C1 enzyme (AKR1C1), NAD(P)H quinone oxidoreductase 1 (NQO-1) were analysed. BPGA exposure resulted in an increase in fold change of NFE2L2 compared to the control, with values changing from 1.00 in control cells to 0.99±0.3 at 3 mM, 1.08±0.04 at 6 mM, and 1.92±0.5 at 9 mM BPGA (ANOVA, p=0.021), as shown in Figure 2C(i). BPGA significantly induced HMOX-1 expression (∼4.7±1.2-fold increase) at 9 mM (ANOVA, P < 0.001) compared to control cells (untreated) (Figure 2C(ii)). This robust increase in HMOX-1 suggests a possible oxidative stress response and activation of cytoprotective pathways, consistent with epithelial-mesenchymal transition (EMT) and cellular adaptation to stress (Singhabahu et al., 2023). Nonetheless, lower concentrations of BPGA did not substantially influence HMOX-1 expression. Additionally, AKR1C1, a crucial enzyme in detoxifying reactive aldehydes and preventing oxidative DNA damage, demonstrated dose-dependent upregulation, albeit without attaining statistical significance at any tested BPGA concentration (Figure 2C(iii)). The mean fold change of AKR1C1 was increased from 1.0 in controls to 1.6±0.8 (3 mM), 2.6±0.9(6 mM), and 3 ± 1.4 for 9 mM BPGA-treated cells. NQO-1, another enzyme that contributes to the antioxidant response by detoxifying reactive quinones and protecting cells from damage, was not significantly altered during BPGA exposure in the tested concentrations (Figure 2C(iv)). The BPGA-induced elevation of ROS contributed to upregulation of the HMOX-1 gene, indicating activation of an HO-1-associated antioxidant defence pathway in BPGA-treated A549 cells. To validate this stress response, immunocytochemistry (ICC) of HO-1 was performed. BPGA treatment increased HO-1 fluorescence intensity in A549 cells compared with the control (Figure 2D). ICC analysis demonstrated a significant increase in the MFI of HO-1 at 3mM BPGA (26.9±2.4) compared to the control (10.8±1.7). However, MFI decreased to 16.1±5.6 for 6mM and 13.6±3.9 for 9mM BPGA, indicating that the significant increase in the HMOX-1 mRNA upregulation did not correspond to proportional HO-1 accumulation at higher BPGA concentrations.

### 3.3. BPGA interfere with cell mitochondrial health and alters metabolism

The elevated levels of ROS can lead to mitochondrial membrane depolarisation, lysosomal membrane permeabilisation and metabolic disruption(Tajeddine, 2016). Therefore, to evaluate the potential impact of BPGA treatment on mitochondrial health and cellular metabolism, TMRM assay, Seahorse Mitostress assay and acridine orange staining were performed.

The cationic cell-permeant fluorescent dye TMRM selectively accumulates in mitochondria having a negative membrane potential (ΔΨm), and its fluorescent intensity is directly proportional to the ΔΨm (Figure 3A). An increase in fluorescence intensity indicates healthy mitochondria, whereas a decrease in fluorescence intensity indicates depolarisation or damage (Cevallos et al., 2025). BPGA exposure led to a concentration-dependent decrease in ΔΨm. A significant depolarisation of mitochondria was evident from 6 mM of BPGA exposure onwards, which is also confirmed by microscopic images showing loss of mitochondrial integrity (Figure 3B). The mean fluorescent intensity was decreased from 33561 ± 2674 of NC to 29544 ± 387, 28668 ± 2724, 23238 ± 1540, 14715 ± 562, 11760 ± 859, and 10909 ± 353 for 1, 3, 6, 9, 12, and 15 mM BPGA, respectively (Figure 3C). Therefore, these findings demonstrate that BPGA exposure is compromising mitochondrial health and correlates with the excessive generation of ROS and a significant increase in cytotoxicity.

**Figure 3.**
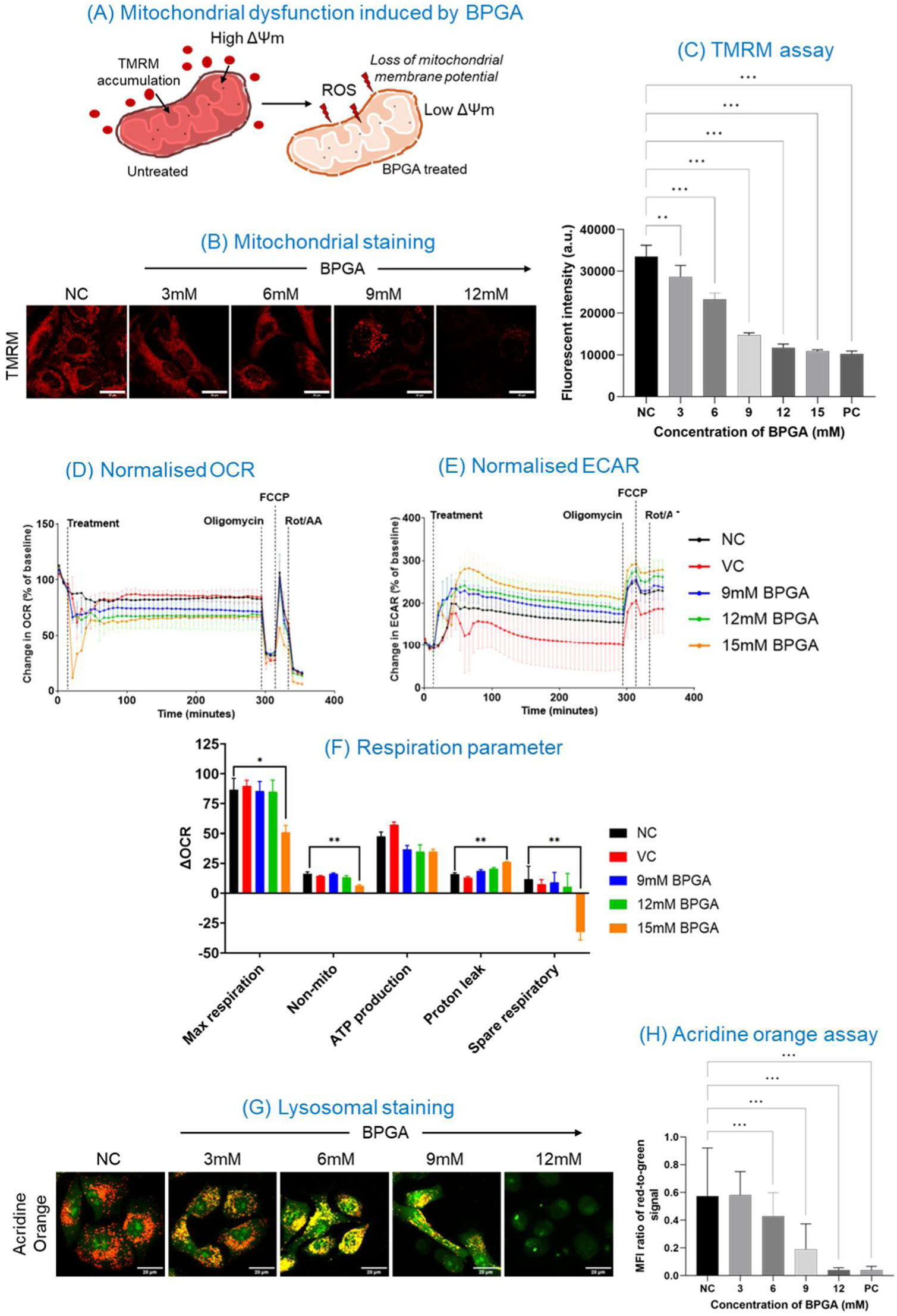
BPGA-induced mitochondrial and lysosomal dysfunction in A549 cells. (A) Schematic representation of mitochondrial impairment upon BPGA exposure. (B-C) Cells were treated with various concentrations of BPGA, 1% phenol H₂O₂ (PC) for 24h and stained with TMRM, and mitochondrial membrane potential was assessed. scale bar represents 20µm. (D) Normalised OCR during the mitochondrial stress test following treatment with BPGA (9–15 mM) or control conditions over a 4-hour period. Sequential injections of oligomycin, FCCP, and Rot/AA were administered to assess key mitochondrial respiration parameters. (E) Normalised ECAR profiles from the same assay, illustrating changes in glycolytic activity over time. (F) Quantitative analysis of respiration parameters derived from the mitochondrial stress test. (G-H) Cells were treated with various concentrations of BPGA, 1% phenol H₂O₂ (PC) for 24h and stained with AO, scale bar represents 20µm. Data are presented as mean ± standard deviation. Statistical comparisons were performed using the ordinary one-way ANOVA test, followed by Dunnett’s multiple comparison test; *p < 0.05, **p < 0.01, ***p < 0.001.

Furthermore, BPGA induced clear, concentration-dependent alterations in mitochondrial respiration and glycolytic activity in A549 cells. Normalised OCR profiles showed a progressive decline in basal respiration with increasing BPGA concentration, becoming apparent at 9 mM (Figure 3D). Most evidently, OCR was reduced by nearly 80% within the first 30 minutes upon 15 mM BPGA exposure. Partial recovery occurred over time, reaching levels similar to 12 mM BPGA. Following FCCP injection, maximal respiration, which represents peak electron transport chain activity, was markedly suppressed in BPGA-treated groups. The inhibition of maximal respiration indicated reduced mitochondrial reserve capacity, while the significant decrease in proton leaks suggested loss of respiratory efficiency associated with impaired coupling (Figure 3F). In contrast, extracellular acidification rate (ECAR) profiles (Figure 3E) showed a pronounced compensatory increase in glycolytic activity. ECAR in A549 cells increased by approximately 2-fold following treatment with BPGA at 9 mM and 12 mM and approached a 3-fold increase at 15 mM, relative to the negative control.

Exposure of BPGA to alveolar epithelial cells led to a significant lysosomal membrane permeabilisation (LMP), as evidenced by acridine orange (AO) staining (Figure 3H). The findings indicate that BPGA compromises lysosomal membrane integrity in a concentration-dependent manner. At the lowest concentration of BPGA exposure (3 mM), A549 cells exhibited distinct, bright red puncta typical of intact lysosomes (Figure 3G), with the red-to-green fluorescence ratio remaining statistically indistinguishable from that of the negative control (untreated cells), indicating a lack of significant LMP. However, at higher concentrations of BPGA exposure (≥ 6 mM) led to a significant decrease (ANOVA, p < 0.001) in the red-to-green MFI ratio. The MFI ratio dropped from 0.574 ± 0.3 (NC) to 0.43 ± 0.1 (6 mM), 0.19 ± 0.1 (9 mM), and 0.04 ± 0.01 (12 mM). The change in the ratio of MFI was also mirrored in live cell imaging, in which a clear transition from the bright red punctate staining in the control group to a diffuse, predominantly green fluorescence in treatment groups was observed (Figure 3G). This decline is due to the loss of the acidic environment within lysosomes, confirming significant LMP.

To check how oxidative stress impacts mitochondria in A549 cells and whether this effect could potentially be reversed using a scavenger molecule, ROS assay and mitochondrial staining assay were performed in the presence of a scavenger molecule (1mM ascorbic acid) (Figure S2 a). The addition of ascorbic acid significantly reduced the BPGA induced ROS generation in A549 cells. Additionally, mitochondria were stained using MitoTracker Orange, a dye whose accumulation and fluorescence are strongly dependent on mitochondrial membrane potential (ΔΨm) (Figure S2b). In parallel, cytosolic oxidative species were assessed using CytoROS Deep Red, a DCFH-based probe that predominantly detects hydrogen peroxide (H₂O₂) following intracellular oxidation, thereby serving as a readout of peroxide-associated oxidative stress.

BPGA induces dose-dependent mitochondrial stress characterised primarily by structural disruption and loss of mitochondrial integrity. In the negative control, mitochondria display elongated, interconnected filamentous networks consistent with preserved ΔΨm and normal bioenergetic status. Upon BPGA exposure, mitochondria become progressively fragmented and swollen, transitioning from organised tubular networks to punctate and aggregated structures. At higher concentrations, diffuse or whole-cell MitoTracker staining is observed, consistent with collapse of ΔΨm and loss of mitochondrial selectivity, indicating substantial mitochondrial dysfunction. Furthermore, the spatial overlap between mitochondrial staining and CytoROS signals supports a mitochondrial origin of oxidative stress. The pattern of localised punctate staining corresponds to mitochondrial hotspots, reinforcing the interpretation that oxidative events are concentrated within damaged mitochondrial networks rather than being uniformly distributed throughout the cytosol. In addition, 1 mM ascorbic acid provides partial protection against BPGA-induced mitochondrial damage. BPGA co-treatment with ascorbic acid partially preserves mitochondrial morphology, with improved network continuity and reduced structural collapse compared to BPGA treatment alone. This protective effect is consistent with ascorbic acid’s ability to scavenge superoxide and hydroxyl radicals, thereby limiting radical-mediated mitochondrial membrane injury and supporting maintenance. Collectively, these findings support the conclusion that BPGA induces a mitochondrial-localised oxidative stress, leading to progressive mitochondrial fragmentation and membrane potential disruption, whereas ascorbic acid partially mitigates radical-driven mitochondrial damage and supports the maintenance of mitochondrial integrity.

### 3.4. BPGA suppresses apoptotic signalling

Excessive intracellular ROS generation can cause mitochondrial stress and activate apoptosis (C.-C. Wu & Bratton, 2013). In the intrinsic apoptotic pathway, ROS induces BCL2-associated X protein (BAX) translocation to the mitochondrial outer membrane, which promotes membrane permeabilisation, leading to pore formation and cytochrome C release into the cytosol. In contrast, B-cell lymphoma 2 protein (BCL-2) counteract BAX, by stabilising the mitochondrial membrane and preventing CYCS release. The free CYCS can bind to Apaf1, driving apoptosome formation and recruiting and activating caspase 9 (CASP9). CASP9 will further activate caspase-3 (CASP3), leading to the execution of apoptosis (Figure 4A). Therefore, to evaluate the effect of BPGA on the progression of apoptosis in A549 cells, the gene expression of various apoptotic-associated proteins (CYCS, CASP9, BAX, BCL-2) were analysed.

**Figure 4.**
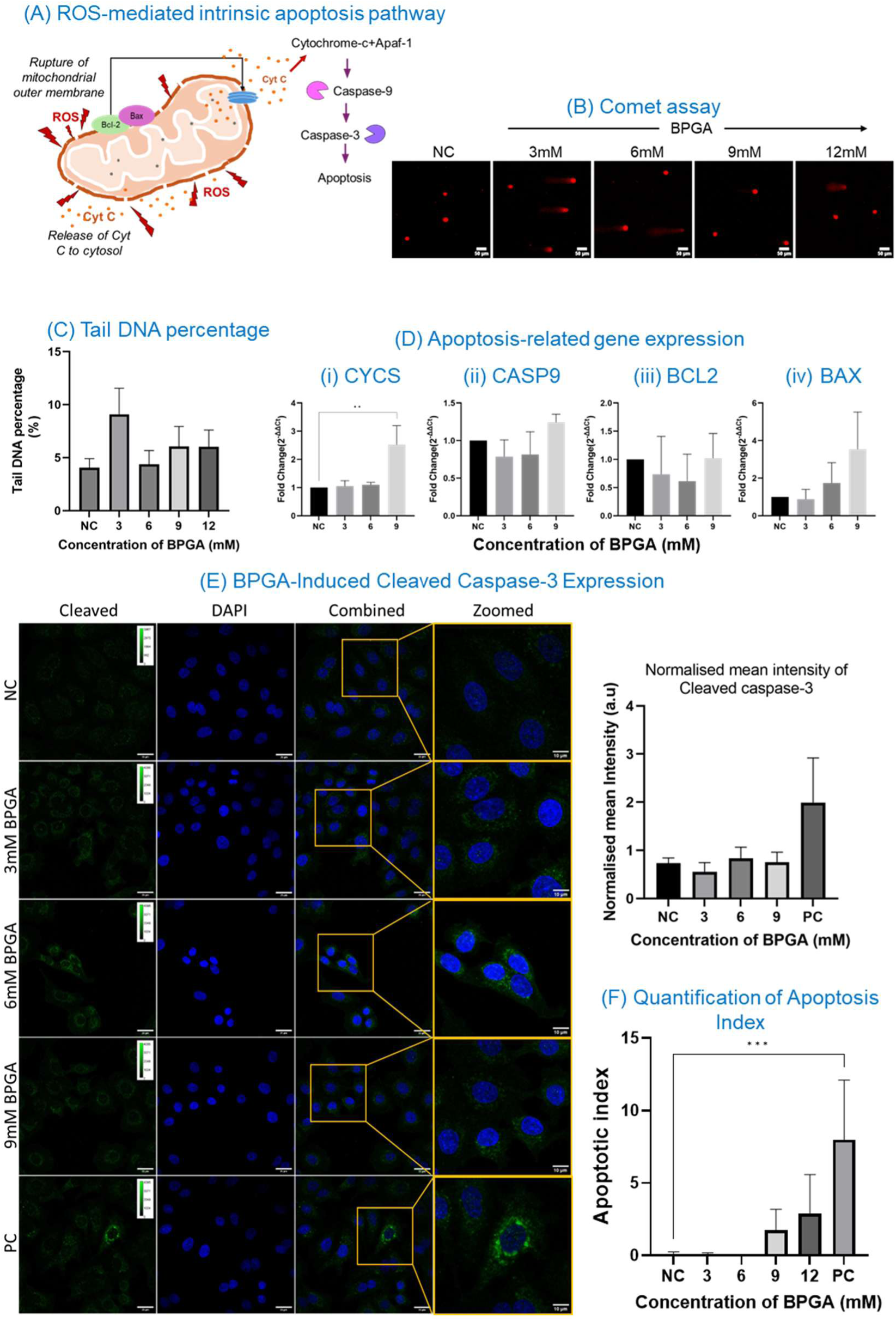
BPGA-induced apoptotic responses in cells. (A) Schematic representation of the ROS-mediated intrinsic apoptosis pathway. (B)Comet assay images showing DNA strand breaks as comet tails after BPGA exposure. (C) Quantification of tail DNA percentage from comet assay performed using OpenComet plugin Fiji-ImageJ. (D) Quantitative PCR analysis of the apoptosis pathway related genes after BPGA treatment, normalised with housekeeping genes (i) CYCS, (ii) CAS9, (iii) BCL2 and (iv) BAX. (E) Cleaved caspase 3 activity on A549 cells treated with various concentrations of BPGA and its normalised mean fluorescence intensity, 25 µM Antimycin A (PC); scale bar represents 25µm (10µm for zoomed images. (F) Quantification of the apoptotic index based on selective uptake of acridine orange (AO) and ethidium bromide (EB) based on membrane integrity.

At low BPGA concentrations (3 mM and 6 mM), CYCS expression was reduced compared to control cells (untreated), indicating suppressed apoptotic signalling or increased stress resistance at lower doses (Figure 4D(i)). However, at higher BPGA concentrations, CYCS expression was significantly increased, suggesting mitochondrial stress and possible activation of the apoptotic pathway. Similarly, the CASP9 showed the same trend in gene expression (Figure 4D(ii)). At lower BPGA concentrations, the fold change in CASP9 was reduced (0.79 ± 0.2 at 3 mM and 0.81 ± 0.3 at 6 mM), whereas at the higher concentration (9 mM), it increased to 1.24 ± 0.1 compared to control (untreated) cells. The BAX gene showed a concentration-dependent increase in fold change relative to control (untreated) cells (0.87 ± 0.5, 1.74 ± 1.0, and 3.54 ± 1.9 at 3 mM, 6 mM, and 9 mM, respectively), indicating activation of the intrinsic apoptotic pathway (Figure 4D(iv)). Nevertheless, the observed elevation in BAX fold change failed to achieve statistical significance. In contrast, BCL2 gene expression showed a minimal change. The fold change of BCL2 is slightly decreased at lower concentrations (3 mM & 6 mM) of BPGA exposure but returned to the baseline at 9 mM (Figure 4D (iii)). This may indicate that lower levels of BPGA exposure reduce anti-apoptotic signals, thereby lowering the threshold for apoptosis activation. Additionally, BPGA-treated cells showed a modest, though not statistically significant, increase in cleaved caspase 3 fluorescence compared with the control (Figure 4E). The normalised mean intensity of cleaved CAS3 was varied from 0.73 ±0.1 for NC to 0.56 ±0.2 (3 mM), 0.83±0.2 (6 mM), 0.75±0.2(9 mM), 1.99 ±0.9 (PC), respectively. The low level of cleaved CAS3 indicates that only a small population of cells undergoes apoptosis, consistent with the qPCR findings.

Apoptosis generally results in DNA fragmentation, membrane blebbing, externalisation of phosphatidylserine and membrane permeabilisation. Therefore, to validate suppressed activation of the apoptotic pathway, single-cell gel electrophoresis (SCGE) or comet assay and AO/EB were conducted. The comet assay detects DNA damage at the single-cell level by quantifying strand breaks which manifest as comet tails under fluorescence microscopy (Figure 4B). In contrast, AO/EB assay can identify the apoptotic cell population through selective uptake of acridine orange (AO) and ethidium bromide (EB) based on membrane integrity. Acridine orange (AO)-permeates both live and dead cells, binds to the DNA and stains it green, however EB enters to the cells with compromised membrane, binds to DNA, and fluoresces a red colour. BPGA treatment elicited a modest elevation in tail DNA percentage, though this alteration lacked statistical significance (Figure 4C). At 3mM of BPGA the percentage of tail DNA was 9.09%±2.4; at 6mM, it was 4.38% ±1.2, at 9mM, 6.04%±1.9 and at 12mM, 6.04%±1.6 in comparison the control (untreated) group showed 4.05%±0.9. The apoptotic index, determined as the ratio of apoptotic cells to total cells, exhibited a concentration-dependent increase; however, this change lacked statistical significance (Figure 4F). At the lower concentration of BPGA exposure, the apoptotic index remained close to zero. A variation was observed beginning at 9mM BPGA, where the apoptotic index was calculated as 1.75±0.6. At 12mM the apoptotic index further increased to 2.88±1.1. Collectively, the results indicate that although BPGA exposure initiates the apoptotic pathway, it does not progress to the complete apoptosis. Only a small population of cells enters apoptosis, and this change is not statistically significant.

### 3.5. BPGA modulates immune response

Because BPGA exposure significantly increases intracellular ROS while suppressing apoptosis, cells may shift toward inflammatory signaling as a possible compensatory response. ROS can activate various redox-sensitive transcription factors such as Nrf2, nuclear factor kappa-light-chain-enhancer of activated B cells (NF-κB) and these molecules can translocate to the nucleus and act as transcription factors for various proinflammatory cytokines (Buelna-Chontal & Zazueta, 2013) and such as tumour necrosis factor-alpha (TNF-α), interleukin-6 (IL-6) and transforming growth factor-beta (TGF-β) (Figure 5A). Therefore, to study the possible changes in the inflammatory mediators’ gene expression associated with the BPGA exposure, the fold change of IL-6, NF-κB, TNF-α, and TGF-β was evaluated (Figure 5B). The gene expression studies revealed a dose-dependent increase in fold changes for IL-6, NF-κB, TNF-α and TGF-β. The fold change of NF-κB was almost similar in 3 mM (1.50 ± 0.8) and 6 mM (1.41 ± 0.3) BPGA-treated groups, while it significantly increased to 2.38 ± 0.06 (ANOVA, P < 0.05) (Figure 5B (i)). Similarly, the fold change in TNF-α gene expression exhibited a progressive, statistically significant increase to 1.71 ± 0.8 for 3 mM, 2.15 ± 0.9 for 6 mM, and 4.02 ± 1.8 (ANOVA, P < 0.05) for 9 mM BPGA-treated groups (Figure 5B(ii)). IL-6 gene expression exhibited a statistically significant, pronounced elevation in fold change(ANOVA, P < 0.001) compared with the control (untreated) cells (Figure 5B(iii)). The fold change of IL-6 was increased to 1.50 ± 0.2 for 3 mM, 3.15 ± 0.3 for 6 mM, and 33.9 ± 6.3 for 9 mM BPGA-treated cells compared with the control. Further, the fold change associated with TGF-β upon the BPGA exposure was increased to 1.01 ± 0.5 for the 3 mM treated group, 1.74 ± 0.2 for the 6 mM treated group, and 2.76 ± 1.1 (ANOVA, P < 0.05) for the 9 mM BPGA treated group (Figure 5B(iv)).

**Figure 5.**
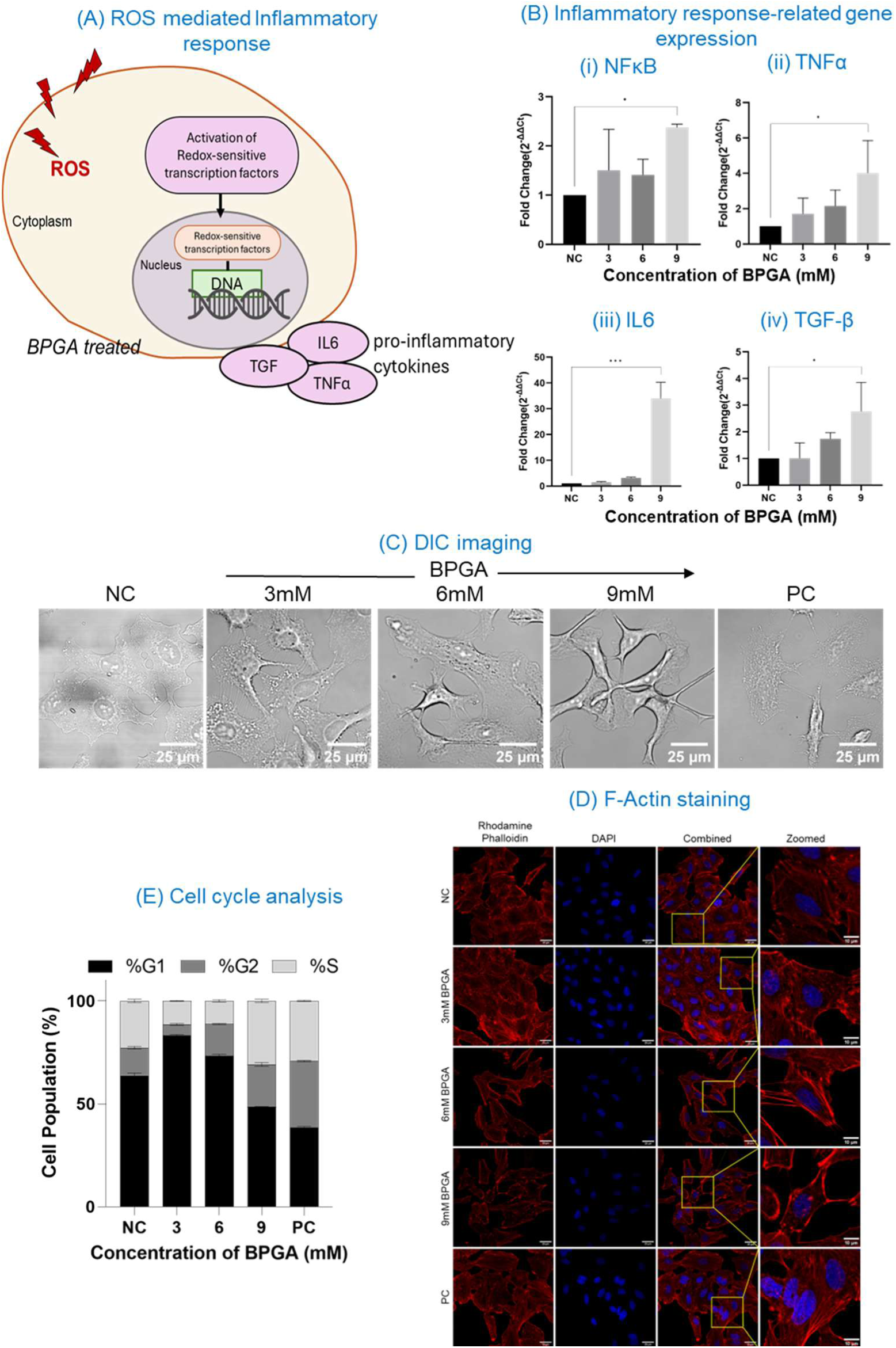
Effects of reactive oxygen species (ROS) on cellular inflammation, morphology, cytoskeletal organization, and cell cycle progression. (A) ROS-mediated inflammatory response. (B) Inflammatory response-related gene expression after BPGA treatment, normalised with housekeeping genes (i) NFκB, (ii)TNFα, (iii) IL6, and (iv) TGF-β, demonstrating differential regulation under ROS exposure. (C) DIC imaging: Representative differential interference contrast (DIC) micrographs illustrating morphological changes in cells subjected to BPGA treatment (scale bar = 25 µm f BPGA, 20ng/mL TGF-β (PC). (D) Phalloidin-Rhodamine/DAPI staining showing the cytoskeleton of A549 cells treated with various concentrations of BPGA, 20ng/mL TGF-β (PC). scale bar represents 25µm (10µm for zoomed). (E) Cell cycle analysis: Distribution of cells across G0/G1, S, and G2/M phases under different BPGA exposure conditions, 20µg/mL mitomycin (PC), highlighting shifts in cell cycle progression.

### 3.6. BPGA induces EMT-Like changes as an adaptive stress response

BPGA treatment of A549 cells elicited a concentration-dependent augmentation of intracellular ROS production and inflammatory responses, concomitant with suppression of apoptosis. The combination of elevated ROS levels and enhanced inflammatory signalling may trigger EMT-like changes in epithelial cells (Xu et al., 2025). The EMT changes in epithelial cells are generally characterised by spindle-shaped morphology, increased expression of mesenchymal markers, enhanced extracellular matrix deposition and contractility and decreased expression of epithelial markers.

BPGA exposure induced a concentration-dependent alterations in cellular morphology, characterised by an elongated, spindle-shaped appearance compared with normal (Figure 5C). Additionally, Immunofluorescence staining of actin filaments using rhodamine phalloidin revealed distinct, concentration-dependent, progressive morphological alterations in A549 cells treated with BPGA (Figure 5D), as observed in DIC imaging. In untreated cells, indicating a normal cobblestone morphology. However, upon BPGA exposure, cytoskeletal rearrangement occurred with noticeable alterations in fibre thickness and structure, especially from 6 mM BPGA onwards. Cells with 9 mM BPGA exposure exhibited denser actin filaments and spread out more throughout the cytoplasm. These results clearly indicate that BPGA exposure triggers significant cytoskeletal remodelling in A549 cells. Also, this remodelling promotes cellular plasticity and migratory behaviour-markers of EMT.

During EMT, epithelial cells often undergo growth arrest, most commonly at G1/S phase (Hu et al., 2025). This arrest will facilitate the transcriptional reprogramming and cellular plasticity, preconditioning cells for EMT. To determine whether BPGA exposure modulates cell cycle distribution in A549 cells, flow cytometric analysis was conducted (Figure 5E). In untreated cells, the cell cycle profile exhibited a balanced distribution typical of a proliferating population, with 63.6% ±0.8 in G0/G1, 22.8%±0.5 in S, and 13.6%±0.4 in G2/M phase. However, at 3mM of BPGA exposure, a strong G0/G1 arrest was induced with 83.3%±0.3 of the cell population in the G0/G1 phase, while the S and G2/M phase was reduced to 11.5%±0.2 and 5.26%±0.4, respectively. This suggests that low doses of BPGA drive cells to a quiescent-like state, consistent with EMT initiation. Further at 6mM BPGA exposure, the G0/G1 phase remained dominant with a cell population of 73.4%±0.6 but G2/M phase rose up to 15.4%±0.2 with 11.3%±0.5 cells in S phase, indicating a mixed population, partially relieving the arrest and reflecting a heterogeneity in EMT progression. At 9mM of BPGA treatment, G0/G1 fraction dropped sharply to 48.5% ±0.2, while S phase and G2/M phase rose to 30.9%±0.7 and 20.6%±0.9 respectively (Figure S3). This redistribution suggests that the cells are no longer arrested and are cycling with altered distribution. Coupled with the observed cytoskeletal alterations and morphological changes, these findings imply that the epithelial cells have reprogrammed and re-entered the cell cycle with mesenchymal traits.

BPGA exposure led to characteristic morphological and biophysical changes, as shown by long-term holotomographic imaging over 48 hours (Figure 6B) and normalised analysis (Figure S4) relative to the initial cycle of each experiment. In the initial 20 hours, both NC and VC maintained a compact, squamous epithelial alveolar type-II phenotype, with confluence remaining above 95% throughout (Figure S4a). In contrast, 9 mM BPGA treatment led to a mild decline in confluency to approximately 90%, accompanied by a 12–15% increase in mean surface area and a transient rise in eccentricity to 120% of baseline, indicating mild elongation and early EMT-like changes. At 12 mM BPGA, cell confluence decreased further to 70%, and the mean surface area reached 135% of the NC values, with a visible gap between cells and an irregular shape. These cells also exhibited increased dry mass (∼20% above NC) and granularity (over 40%), further reflecting cytoskeletal remodelling and increased intracellular density, both of which are linked to a more active metabolism. By comparison, cells exposed to 20 ng/mL TGF-β, a known positive EMT control, showed similar morphological changes, with a 20% decrease in confluency and a 30% increase in mean surface area, further confirming EMT induction. Further, live-cell imaging from 24 to 48 hours showed continued but stabilised structural remodelling following BPGA exposure (Figure S4b). The BPGA treatment resulted in a marginal change in confluence, with 9 mM stabilising at ∼98% and 12 mM at ∼102%. However, the mean surface area increased in BPGA-treated cells, particularly at 12 mM, reaching 25% above that of NC. In addition, eccentricity remained stable across all treatment groups, suggesting that elongation was limited to the initial exposure period. The mean dry mass varied in different ways: NC showed a decline and recovery after 35 hours; VC remained near baseline; 9 mM BPGA slightly decreased (92%); and 12 mM BPGA continued to rise to 118%, indicating sustained intracellular accumulation. In contrast, the granularity showed the strongest treatment effect, increasing by up to ∼120% in the 9 mM group and ∼180% in the 12 mM BPGA group, whereas NC and VC remained near baseline.

**Figure 6.**
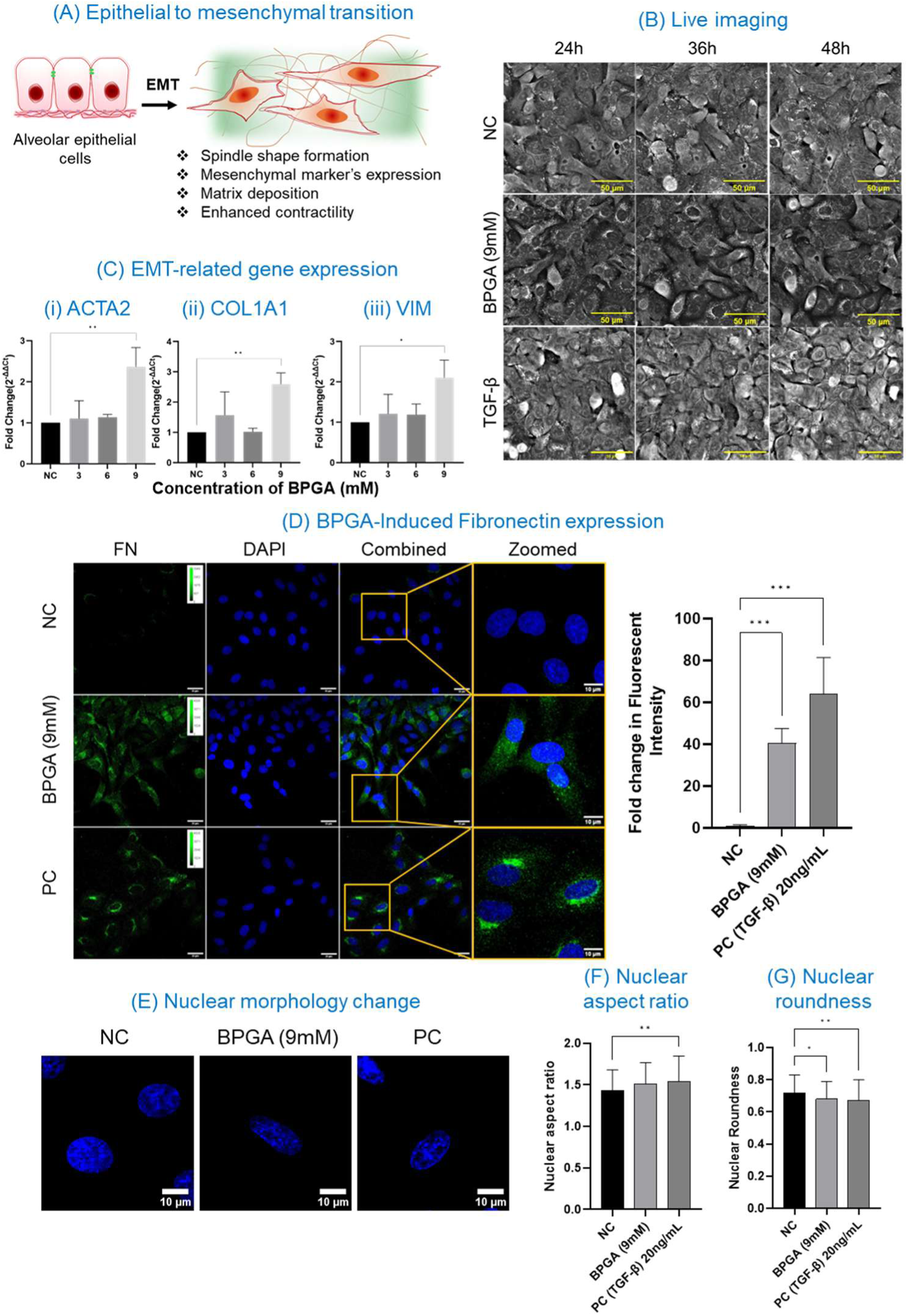
BPGA-induced epithelial to mesenchymal transition (EMT) and associated cellular changes. (A) Schematic representation of EMT. (B) Holotomographic images at 24, 36 and 48 h show that 9 and 12 mM BPGA treatments caused progressive cell elongation, intercellular gaps, and irregular morphology compared with NC and VC. TGF-β (20 ng/mL) induced moderate elongation without marked loss of compactness. Scale bars: 50 μm. (C) Quantitative PCR analysis of the EMT-related genes after BPGA treatment, normalised with housekeeping genes (i)ACTA2, (ii) FN, (iii) VIM. (D) FN expression and quantification of FN fluorescence intensity in A549 cells treated with BPGA (9mM), PC-TGFβ (20ng/mL) and NC-Media alone for 24, scale bar represents 25µm (10µm for zoomed images). Quantification of FN fluorescence intensity expressed as fold change relative to NC (n = 5). (E) Representative confocal images demonstrating the changes in the nuclear morphology, (F)Quantification of nuclear aspect ratio, (G) Quantification of nuclear roundness. Statistical comparisons were performed using the one-way ANOVA, and Dunnett multiple comparison test versus the normal control (NC); *p < 0.05, **p < 0.01, ***p < 0.001.

**Figure 7.**
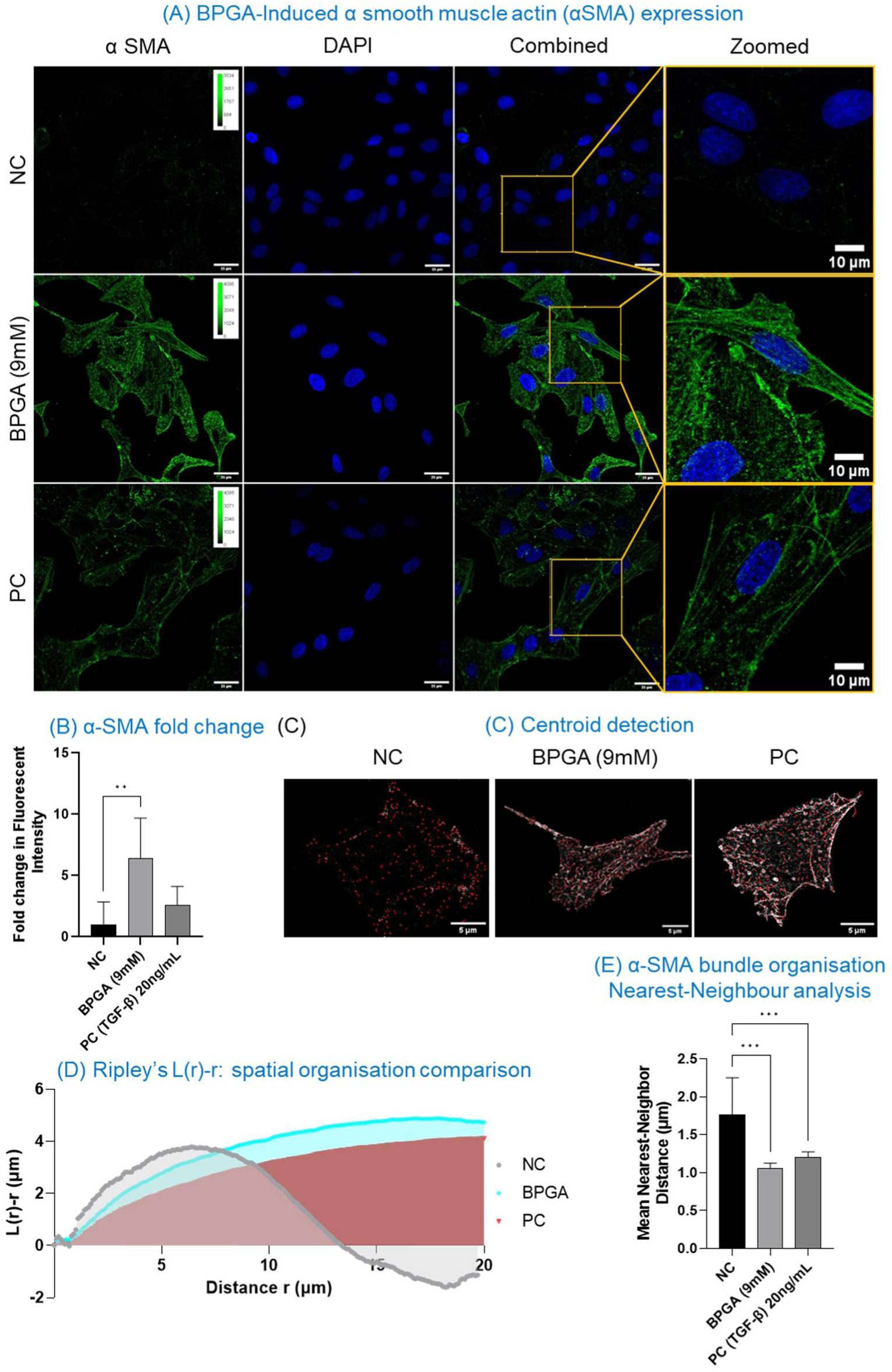
Analysis of α-SMA Expression and Spatial Organisation in A549 Cells. (A) α-SMA expression in A549 cells treated with BPGA (9mM), PC-TGFβ (20ng/mL) and NC-Media Alone for 24h, scale bar represents 25µm (10µm for zoomed insets(B) Quantification of α-SMA fluorescence intensity expressed as fold change relative to NC (n = 5). (C) Representative confocal image of individual cells showing the centroid overlays using the ThunderSTORM plugin, (D) Spatial organisation comparison of α-SMA using Ripley’s L(r)-r function, (E) Nearest neighbour analysis of α-SMA bundle organisation across treatment groups. Statistical comparisons were performed using the one-way ANOVA, and Dunnett multiple comparison test versus the normal control (NC); *p < 0.05, **p < 0.01, ***p < 0.001.

During EMT, a significant increase of mesenchymal markers can be observed both transcriptionally and translationally, along with a clear morphology change (Figure 6A). Therefore, to validate the acquisition of mesenchymal characteristics, gene expression of EMT-related markers, including smooth muscle alpha actin (ACTA2), vimentin (VIM) and collagen type I alpha 1 chain (COL1A1) was studied. BPGA exposure significantly increased the fold change of ACTA2, COL1A1, and VIM, specifically at 9 mM in the BPGA treatment groups (Figure 6C). The BPGA exposure led to 1.1 ± 0.4-fold change in ACTA2 at 3 mM, which further increased to 1.13 ± 0.06 at 6 mM and 2.36 ± 0.5 at 9 mM (ANOVA, adjusted p-value 0.002) (Figure 6C (i)). Similarly, for COL1A1, BPGA exposure resulted in a fold change of 1.57 ± 0.8 in the 3 mM-treated group, 1.02 ± 0.1 at 6 mM, and 2.59 ± 0.3 at 9 mM (ANOVA, adjusted p-value 0.005) (Figure 6C (ii)). Additionally, a fold change of 1.21 ± 0.5 was observed for VIM in the 3 mM BPGA-treated group, 1.19 ± 0.3 at 6 mM, and 2.10 ± 0.4 at 9 mM (ANOVA, adjusted p-value 0.013) (Figure 6C(iii))

Additionally, to validate the transcriptional upregulation of mesenchymal markers observed by qPCR, immunocytochemistry was performed to assess cytoplasmic α-SMA expression, and fibronectin (FN) deposition was analysed. Fibronectin is an extracellular matrix glycoprotein that is upregulated during injury, inflammation, and epithelial-to-mesenchymal transition. An Increased fold change in mean fluorescence intensity of fibronectin (Fold change of 40.5 ± 7.1, ordinary one-way ANOVA, adjusted p-value< 0.001) (Figure 6D) was observed in BPGA-treated (9mM) cells compared to the control (untreated). Concurrently, a variation of nuclear morphology was also found in the BPGA-treated group (9mM) (Figure 6E). Briefly, the nuclei of BPGA-treated cells exhibited an increased aspect ratio (1.51 ± 0.2) compared to untreated cells (1.43 ± 0.2) (Figure 6F). The roundness of the nuclei was statistically decreased upon BPGA treatment (9mM) (0.681 ± 0.1) in comparison with the untreated cells (0.718 ± 0.1, ordinary one-way ANOVA, adjusted p-value 0.031) (Figure 6G).

During EMT, epithelial cells lose polarity and adhesion. α-SMA integrates into stress fibres, conferring on cells the contractile properties needed for migration. This structural change is a hallmark of mesenchymal identity. α-SMA is widely recognised as a myofibroblast marker, indicating that epithelial cells have transitioned into a phenotype capable of producing extracellular matrix and exerting mechanical force. Transforming growth factor-β (TGF-β), a principal inducer of EMT, directly upregulates α-SMA expression, making α-SMA a downstream readout of canonical EMT signalling. BPGA treatment significantly increased α-SMA staining (Fold change of 6.07± 1.8 (Figure 9B), Ordinary one-way ANOVA, adjusted p-value 0.003), with prominent fibre-like structures organised into filamentous bundles, characteristic of mesenchymal/myofibroblast phenotypes. Spatial distribution analysis revealed that BPGA treatment not only increases α-SMA expression but also alters its organisation into distinct bundles. Ripley’s L(r) – r analysis was applied to determine whether α-SMA was randomly distributed, clustered, or dispersed. BPGA-treated cells (9 mM) showed significantly altered α-SMA organisation, with a positive deviation (Ripley’s L(r) – r > 0) compared to untreated cells (Ripley’s L(r) – r close to zero), but similar to TGF-β-treated cells (Ripley’s L(r) – r > 0), indicating non-random, structured assembly of stress fibers. In the BPGA-treated group (9 mM), the L(r) – r value steadily increased, indicating sustained clustering of spatial features, with stronger organisation as distance increased. However, in the negative control, dispersion was observed at larger scales, indicating weak clustering that dissipates with distance. Complementarily, the average spacing between bundles was determined using nearest-neighbour analysis of α-SMA bundles. BPGA treatment (9 mM; NND = 1.05 ± 0.01 µm) reduced bundle spacing, meaning bundles were more tightly organised compared to untreated cells (NND = 1.76 ± 0.5 µm). This reflects enhanced cytoskeletal remodelling and contractile fibre formation, consistent with EMT progression. These findings highlight that BPGA not only upregulates α-SMA expression but also drives its reorganisation into tightly packed stress fibres, reinforcing the acquisition of mesenchymal contractile properties.

## 4. Discussion

Many studies have shown that the thermal degradation of flavoured e-cigarette liquid is the primary contributor to aldehyde formation within e-cigarette vapour (Khlystov & Samburova, 2016), leading to significant concern about user exposure to potentially harmful compounds (Lorkiewicz et al., 2022; Sleiman et al., 2016). Notably, the decomposition products may exhibit distinct toxicological properties in comparison with the parent flavouring molecule (Erythropel et al., 2019), necessitating a comprehensive toxicological evaluation(Ali et al., 2023).

Flavouring agents such as benzaldehyde, vanillin, or cinnamaldehyde present in the e-cigarette liquid react with propylene glycol (PG) through a reversible chemical reaction and form flavour aldehyde PG acetals (Coleman Iii, 2006). Up to 50-80% of this aldehyde PG acetal can be transferred to the e-cigarette vapour and reach the alveolar epithelium. Some e-cigarette liquid contains benzaldehyde as high as 21mg/mL (∼200mM) (Tierney et al., 2016b). Benzaldehyde reacts with propylene glycol (1,2-propanediol) to form geometrical isomers of 4-methyl-1,3-dioxane phenyl collectively known as benzaldehyde PG acetal (BPGA) (Coleman Iii, 2006). The BPGA is known to impair phagocytosis and provoke oxidative burst (Hickman et al., 2019). Additionally, PG acetals can interfere with the various key proteins essential in glycolysis, the pentose phosphate pathway or the electron transport chain (ETC), subsequently altering the normal metabolic function of the cell (Hickman et al., 2019).

While various investigations have been conducted to elucidate the potential toxicity effects of primary e-cigarette liquid components (Al-Saleh et al., 2020; Ebersole et al., 2020; Effah et al., 2022; Phillips et al., 2017). However, investigations into the biological effects of secondary byproducts formed during e-cigarette aerosolisation, such as BPGA, remain limited. Nonetheless, most studies have focused on endpoint cytotoxicity experiments, such as the MTT assay, but it is recognised that sub-lethal concentrations of chemicals can activate various pathways involved in cytotoxicity (Fulda et al., 2010), inflammation(Afolabi et al., 2019), and stress responses, even in the absence of overt cell death (Lee et al., 2023). The use of flavour additives increases the sensory appeal of vaping (Pullicin et al., 2019), thereby making it more pleasant and attractive to users (Goldenson et al., 2016). However, the continued use of such chemicals, linked to longer vaping sessions (Leventhal et al., 2019), ultimately increases the duration of total exposure to potentially harmful chemicals. This prolonged exposure to vaping chemicals can aggravate the risk of cellular damage.

The present study indicated that exposure to BPGA exceeding 9 mM caused considerable cell death, whereas it exhibited minimal or negligible cytotoxicity at concentrations below 9 mM. The cytotoxicity of increased BPGA concentration was demonstrated by metabolic activity (MTT assay) and lysosomal integrity (NR-uptake assay), suggesting that BPGA impairs cell viability via multiple stress mechanisms. The observed cytotoxicity threshold of BPGA corresponds with the cytotoxicity of flavourants in e-cigarette aerosols, such as the significant cytotoxicity (LD50) observed in A549 cells exposed to cherry-flavoured e-cigarette liquid (Findlay-Greene et al., 2025).

BPGA exposure elicited a significant elevation in intracellular ROS production beginning at 6 mM, a concentration that did not reduce cell viability. It confirms that BPGA can increase oxidative stress in alveolar epithelial cells in vitro (Sussan et al., 2015); furthermore, this effect is more pronounced at higher BPGA concentrations. Mechanistically, BPGA at higher concentrations reduces the oxygen consumption rate (OCR), impairs mitochondrial respiration (Kruppa et al., 2018), and causes electron leakage in the mitochondrial electron transport chain. Therefore, the disruption of redox balance eventually increases ROS generation (Jabba et al., 2020), surpassing cells’ antioxidant defence capacity and leading to cellular damage. However, BPGA at sublethal concentration triggered a compensatory antioxidant response, suggesting that cells effectively neutralise ROS-generated stress. To validate this, gene expression of various proteins involved in the antioxidant pathway showed a significant increase in the fold change of heme oxygenase-1 (encoded by HMOX1) compared to NQO-1 or AK1R1, indicating its hierarchical role in the ROS-mediated oxidative stress response (Consoli et al., 2021). The immunocytochemistry assay confirmed increased HO-1 fluorescence in BPGA-treated A549 cells compared with the negative control. The significant increase in HO-1 expression is consistent with its role as the primary stress-responsive enzyme (Consoli et al., 2021) and “gatekeeper” of cell fate decisions under oxidative stress conditions (Raghunandan et al., 2021). However, the HMOX1 promoter contains several stress-responsive elements recognised by various transcription factors, including AP-1, NF-κB, HIF, and STAT, as well as Nrf2 (Gozzelino et al., 2010; Medina et al., 2020). Therefore, this multifactorial regulation explains a significant fold change in HMOX1 despite minimal change in Nrf2 levels, as the downstream target gene exhibits greater expression than its regulatory transcription factors. Nonetheless, at higher BPGA concentrations, HO-1 expression was reduced but remained higher than in untreated cells, indicating impaired regulation under high oxidative stress.

Additionally, the variation in the ECAR profile suggests a metabolic shift towards glycolysis, possibly as a compensatory response to impaired mitochondrial function. The elevation in ECAR persisted throughout the assay, suggesting a sustained dependency on glycolytic metabolism. Quantitative analysis of mitochondrial parameters supported these findings. BPGA exposure significantly reduced non-mitochondrial respiration and spare respiratory capacity, accompanied by a moderate decline in ATP-linked respiration. The marked reduction in spare respiratory capacity indicated impaired mitochondrial adaptability and diminished bioenergetic flexibility in A549 cells under BPGA stress. Therefore, BPGA disrupts mitochondrial function and drives metabolic reprogramming from oxidative phosphorylation toward glycolysis, consistent with early epithelial stress and EMT-like metabolic adaptation. The excessive oxidative stress can damage the lysosomal membrane integrity and further lead to significant lysosomal membrane permeabilisation (Qi et al., 2024). The BPGA-induced LMP can cause lysosomal contents to leak out and subsequently trigger an inflammatory response (Zhang et al., 2023) and cell death signalling (Wang et al., 2018). The lysosomal acidic environment protonates the acetal oxygen, enhances the leaving ability of oxygen and leads to the loss of the attached alcohol group to form a carbocation intermediate, which further accelerates the hydrolysis, resulting in the formation of the corresponding aldehyde (B. Liu & Thayumanavan, 2017). BPGA include a 1,3-dioxolane ring, formed by the reaction between benzaldehyde and propylene glycol, which contains two oxygen atoms characteristic of acetals. Protonation will lead to the breaking of the acetal bond during hydrolysis, releasing benzaldehyde. The reactive benzaldehyde thus formed will further amplify oxidative stress and promote lipid peroxidation, creating a positive feedback loop that further exacerbates lysosomal membrane permeabilization (Hajdú et al., 2023).

Elevated oxidative stress can trigger the intrinsic apoptosis pathway, eventually leading to cell death. The present study demonstrated a significant upregulation of cytochrome C (CYCS) at 9 mM BPGA. In contrast, the mRNA expression levels of caspase 9, Bax or Bcl2 did not show statistically significant changes. ROS are known to activate the proapoptotic proteins Bax and Bak, promoting their oligomerisation and ultimately permeabilising the mitochondrial outer membrane. Mitochondrial outer membrane permeabilisation (MOMP) subsequently triggers cytochrome c release into the cytosol, promoting Apaf-1 oligomerisation and apoptosome assembly, thereby activating the intrinsic caspase cascade, notably caspase-9, to carry out apoptosis. Active caspase-9 proteolytically activates procaspase-3 by cleavage, yielding the mature executioner caspase-3. Subsequently, it cleaves key substrates, including PARP and ICAD, leading to apoptosis. However, the cleaved caspase-3 fluorescence only showed a modest increase in the BPGA-treated A549 cells. Similarly, the comet assay and AO/EB assay did not reveal any significant changes in tail DNA% or apoptotic index, respectively. Collectively, these findings suggest that A549 cells may undergo an adaptive response at sublethal concentrations of BPGA, such that, although mitochondrial stress signals are initiated, they fail to progress to the full apoptotic cascade, allowing cells to evade apoptosis and maintain survival under stressful conditions. It is plausible that once a critical threshold (over 12 mM BPGA) is exceeded, this compensatory mechanism becomes overwhelmed, causing permanent damage to mitochondria.

A549 cells are attempting to develop adaptive mechanisms to survive BPGA exposure at sublethal concentrations through various stress-responsive pathways. Additionally, the gene expression studies of inflammatory markers have revealed a close interconnection between oxidative stress and inflammatory response. BPGA exposure increased ROS levels in a concentreation-dependent manner in A549 cells, activating redox-sensitive transcription factors, including NF-κB, which ultimately upregulated the expression of IL-6, TNF-α, and TGF-β, as observed in many studies. The initial phases of oxidative stress, in which ROS trigger NFκB activation through upstream kinase signalling, promote the transcription of inflammatory cytokines, aligning with the concentration-dependent increase in the relative expression of proinflammatory markers observed at sublethal exposure to BPGA treatment. In contrast, under excessive ROS production, significant cell death was observed, validating the bidirectional role of NFkB signalling in gene regulation. Concurrently, the elevated ROS levels (Richter & Kietzmann, 2016) and LMP activate the various profibrotic pathways (Tian et al., 2025). The sustained inflammation through the oxidative stress–lysosomal dysfunction axis perpetuates fibrotic remodelling. Under sublethal exposure to BPGA, cells remain metabolically active; however, due to the combined effects of oxidative stress and inflammatory signals, epithelial-to-mesenchymal transition (EMT) can be initiated. As an initiation to EMT, the cell arrest at G1/G0 phase was observed at 9mM of BPGA. This arrest will help cells to adapt and precondition themselves to EMT.

Oxidative stress serves as a well-established driver of EMT, primarily through redox-sensitive transcription factors such as NF-κB and cytokine activation, including TGF-β. The BPGA exposure resulted in the significant upregulation of TGF-β; concurrently, the mRNA levels of mesenchymal markers, ASMA, VIM, and COL1A1, canonical for the acquisition of mesenchymal traits, motility, invasiveness, and resistance to apoptosis, were also significantly upregulated in a concentration-dependent manner. Additionally, long-term holotomographic imaging and immunofluorescence staining revealed the acquisition of a fibrotic-type morphology, increased fluorescence intensity of alpha-smooth muscle actin (αSMA) and fibronectin (FN), cellular cytoskeletal reorganisation, αSMA bundle formation, and changes in nuclear morphology upon BPGA exposure. Altogether, it suggests that BPGA, at sublethal concentrations, favours the EMT pathway rather than apoptosis as an adaptive survival mechanism.

## 5. Conclusion

E-cigarettes have gained recognition as a tool for the cessation of conventional smoking, and their use has increased in various age groups. A wide range of flavouring agents is added to improve sensory appeal, but these agents can form secondary products with distinct toxicological profiles from their parent molecules. Currently, there are no globally recognised safety standards for flavouring agents in e-liquids that are based on inhalation toxicology. The present study provides a mechanistic model of alveolar injury and EMT risk induced by benzaldehyde propylene glycol acetal (BPGA), a berry-flavoured e-cigarette adduct, in A549 cells. BPGA exposure caused a concentration-dependent decline in cell viability accompanied by morphological changes, mitochondrial dysfunction and lysosomal destabilisation. BPGA exposure resulted in a concentration-dependent increase in ROS levels, initially upregulating antioxidant enzymes at moderate concentrations but overwhelming this response at higher doses. BPGA at sublethal concentrations upregulated mRNA expression of inflammatory markers, creating a pro-inflammatory milieu. Eventually, it led to altered EMT marker expression in A549 cells, implicating BPGA in EMT. These findings collectively highlight the need for comprehensive safety standards regarding the utilisation of flavouring agents in e-cigarettes. More comprehensive investigations employing sophisticated organ-on-a-chip and exposure-on-chip models(Bernardino de la Serna et al., 2024; X. Liu et al., 2025) or ultimately animal models, are warranted to delineate the long-term effects of BPGA and analogous compounds. Such evidence will be critical to designing health guidelines regarding the use of flavoured e-cigarettes.

## Acknowledgements

The authors acknowledge the director and head of the Biomedical Technology Wing, Sree Chitra Tirunal Institute for Medical Sciences and Technology, Trivandrum, for their support in providing the infrastructure to carry out this work. We deeply acknowledge Dr PV Mohanan (Late) for the guidance, support, and encouragement in completing the study.

## Author contributions

**JX-**Conception and design of study, Acquisition of data, Analysis and interpretation of data, Drafting the manuscript, **YY**-Acquisition of data, Analysis and interpretation of data, Drafting the manuscript, **BMV**-Acquisition of data, Drafting the manuscript, **ZL**-Drafting the manuscript, Critical revision for important intellectual content, **MKB**-, Acquisition of data, Drafting the manuscript, Critical revision for important intellectual content, **RNS**-Acquisition of data, Analysis and interpretation of data, Critical revision for important intellectual content, **APR**-Conception and design of study, Analysis and interpretation of data, Critical revision for important intellectual content, Final approval of the version to be published, Corresponding. **JBS-** Conception and design of study, Analysis and interpretation of data, Critical revision for important intellectual content, Final approval of the version to be published, Corresponding,

## Data Availability Statement

Data is available upon request from the authors.

## Declaration

Consent for publication-All authors agreed to submit the manuscript to the journal, and the same has been approved by the parent institute.

## 7. Supplementary Data

**Figure S1.**
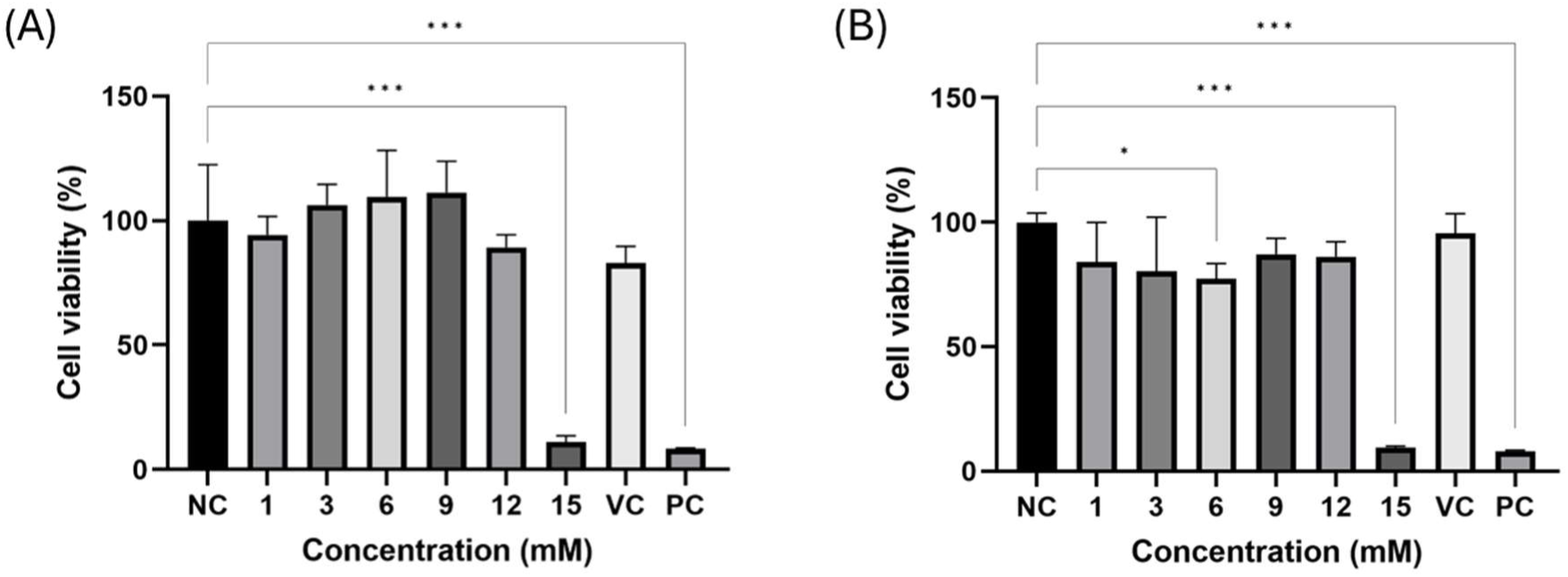
Acute BPGA exposure. Cell viability is expressed by the ability of A549 cells to reduce MTT into formazan crystals. A549 Cells. A549 Cells (0.01 × 10⁶cells/well) were exposed to BPGA (1mM, 3mM, 6mM, 12mM, 15mM) 0.1% DMSO (VC) and 0.1% Phenol (PC) for 2hr (A) and 4hr (B).. Statistical comparisons were performed using the ordinary one-way ANOVA test, followed by Dunnett’s multiple comparison test; *p < 0.05, **p < 0.01, ***p < 0.001

**Figure S2.**
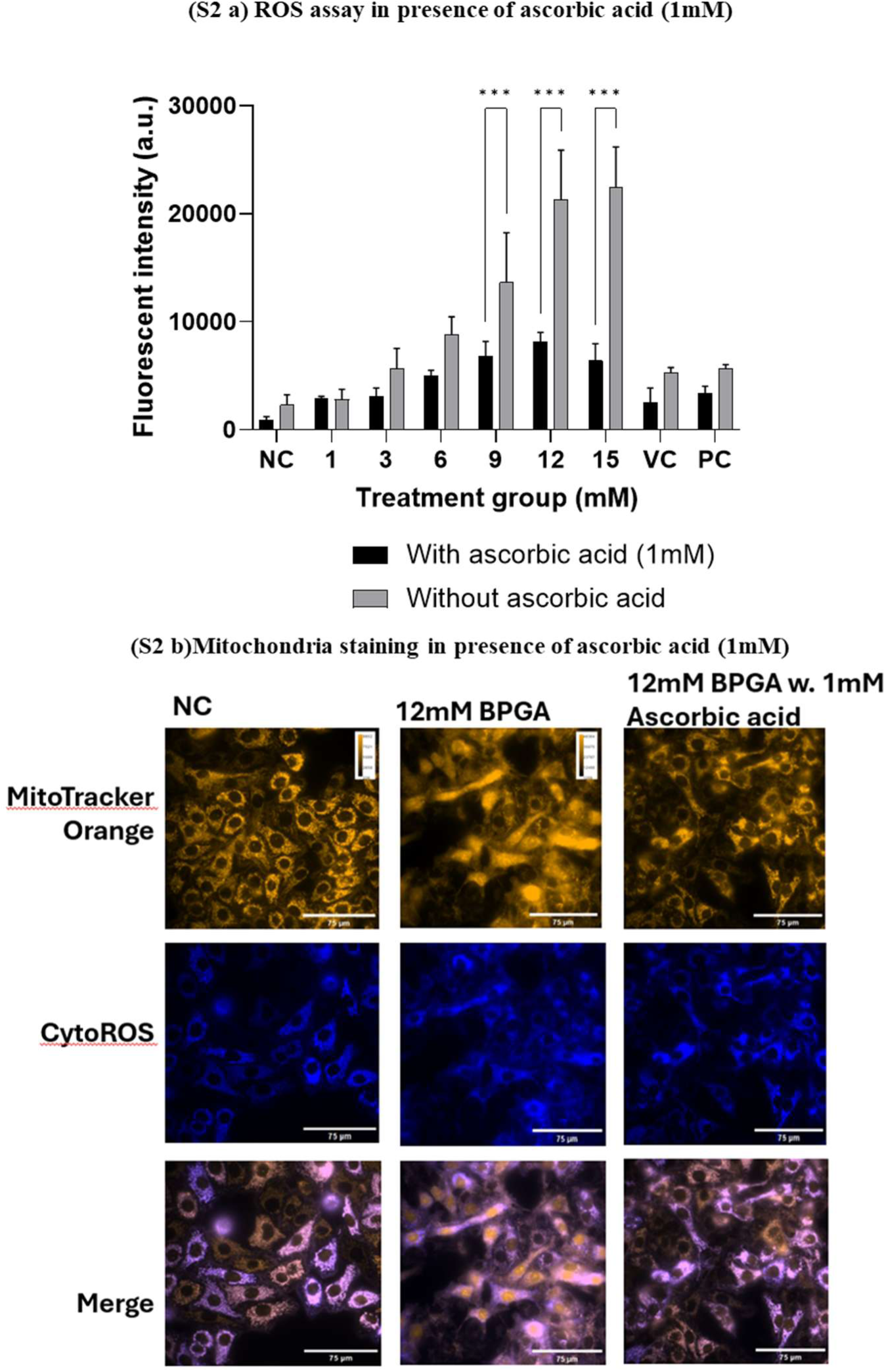
Ascorbic acid partially attenuates BPGA-induced mitochondrial dysfunction. Co-treatment with 1 mM ascorbic acid (AC) protects against BPGA-induced mitochondrial damage, preserving mitochondrial structure compared to BPGA treatment alone.(A)Cells were pretreated with 10µM DCFDA for 30 min. Further cells were treated with various concentrations of BPGA (NC, 3, 6, 9, 12, 15 mM) and 100µM H₂O₂ (PC) for 4h with or without Ascorbic acid. RFI was measured using a multiwell plate reader (Tecan Infinite F Nano+)-excitation/emission filter settings of 488(20)/530(25) nm. (B) A549 cells treated with various concentrations of BPGA (NC, 12 mM) with and without ascorbic acid (AA). Cells were stained with Mito Tracker Orange and CytoROS. Images were captured in a fluorescent microscope.

**Figure S3.**
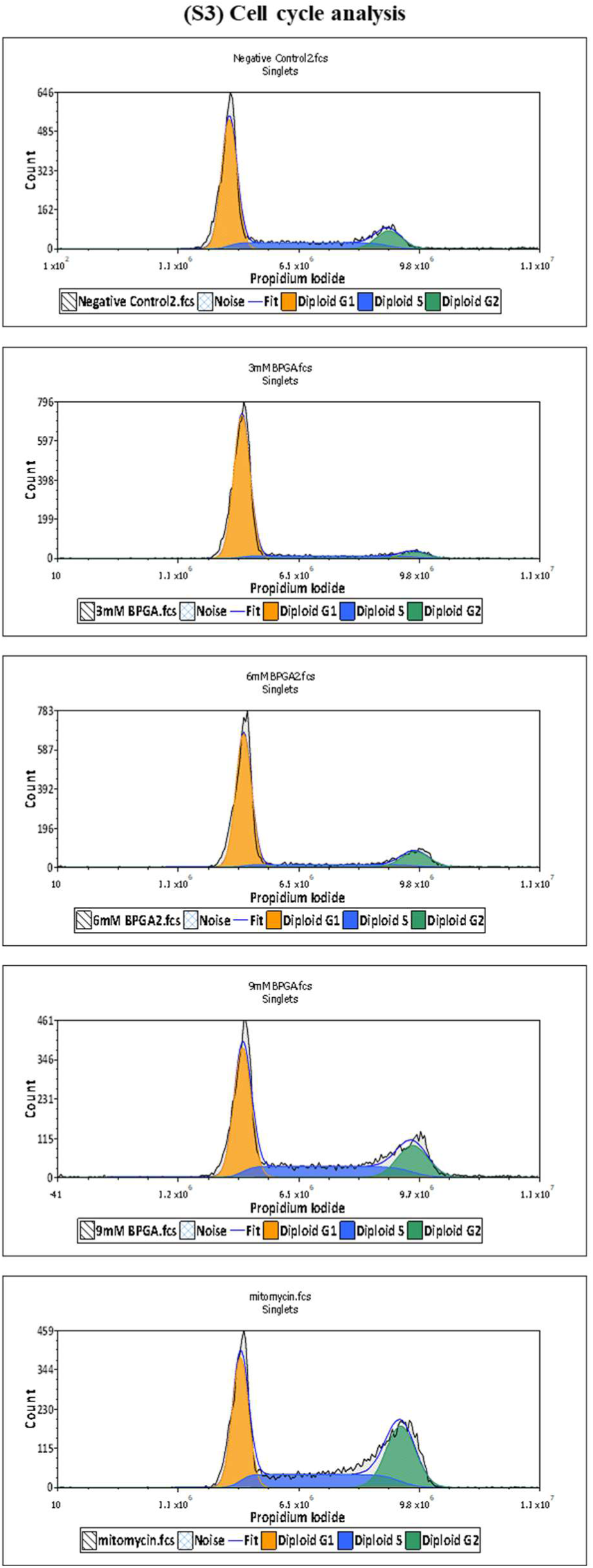
Cell cycle analysis: Distribution of cells across G0/G1, S, and G2/M phases under different BPGA exposure conditions, 20µg/mL mitomycin (PC), highlighting shifts in cell cycle progression.

**Figure S4.**
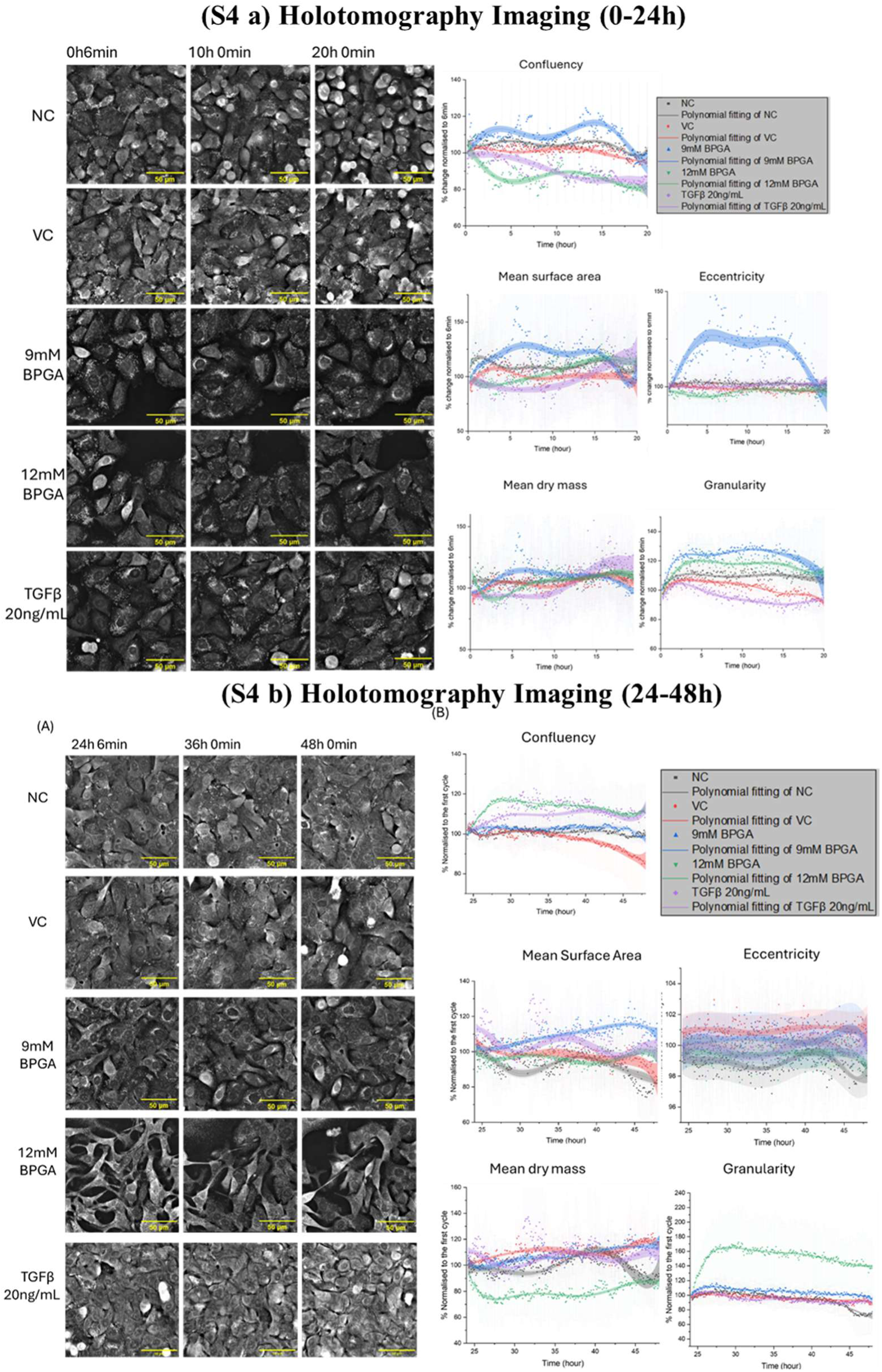
BPGA induces concentration-dependent morphological and biophysical alterations in A549 cells. (A) Holotomographic images at 0,10 and 20 hours, and 24,36,48 hours show that 9 and 12 mM BPGA treatments caused progressive cell elongation, intercellular gaps, and irregular morphology compared with NC and VC. TGF-β (20 ng/mL) induced moderate elongation without marked loss of compactness. Scale bars: 50 μm. (B) Quantitative analysis over 48 h showing changes in confluency, mean surface area, eccentricity, dry mass, and granularity. BPGA increased surface area, dry mass, and granularity in a dose-dependent manner.

**Supplementary Table 1.**
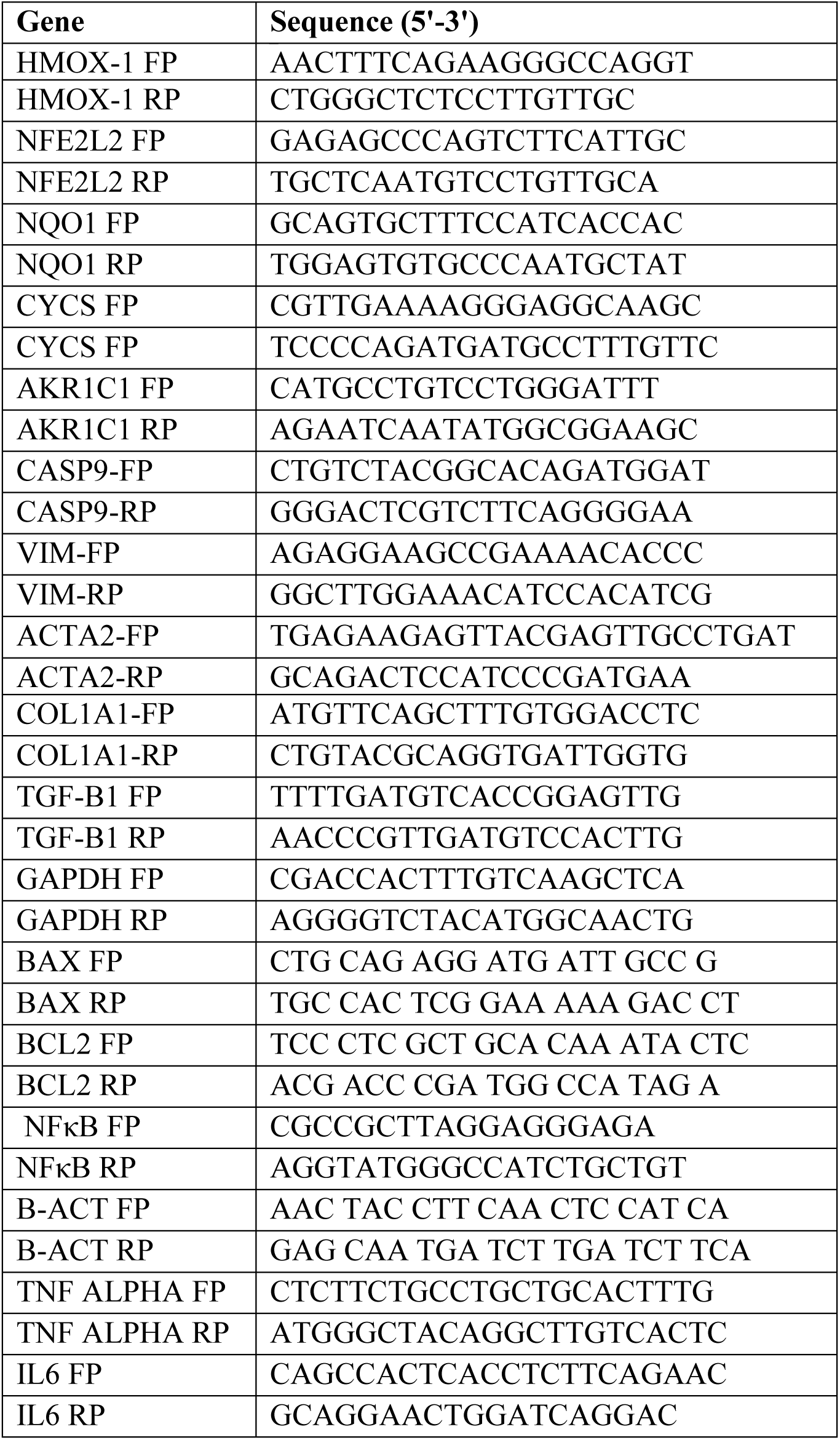
Primers used for qPCR studies.

